# Genome-wide Meta-analysis of 158,000 Individuals of European Ancestry Identifies Three Loci Associated with Chronic Back Pain

**DOI:** 10.1101/244483

**Authors:** Pradeep Suri, Melody R. Palmer, Yakov A. Tsepilov, Maxim B. Freidin, Cindy G. Boer, Michelle S. Yau, Daniel S. Evans, Andrea Gelemanovic, Traci M. Bartz, Maria Nethander, Liubov Arbeeva, Lennart Karssen, Tuhina Neogi, Archie Campbell, Dan Mellstrom, Claes Ohlsson, Lynn M. Marshall, Eric Orwoll, Andre Uitterlinden, Jerome I. Rotter, Gordan Lauc, Bruce M. Psaty, Magnus K Karlsson, Nancy E. Lane, Gail Jarvik, Ozren Polasek, Marc Hochberg, Joanne M. Jordan, Joyce B. J. Van Meurs, Rebecca Jackson, Carrie M. Nielson, Braxton D. Mitchell, Blair H. Smith, Caroline Hayward, Nicholas L. Smith, Yurii S. Aulchenko, Frances M.K. Williams

**Author notes:** Corresponding author: Pradeep Suri, MD, MS VA Puget Sound Health Care System, S-152-ERIC, 1660 S. Columbian Way, Seattle, WA, 98108.

## Abstract

**OBJECTIVES:** To conduct a genome-wide association study (GWAS) meta-analysis of chronic back pain (CBP).

**METHODS:** Adults of European ancestry were included from 16 cohorts in Europe and North America. CBP cases were defined as those reporting back pain present for >3-6 months; non-cases were included as comparisons (“controls”). Each cohort conducted genotyping using commercially available arrays followed by imputation. GWAS used logistic regression models with additive genetic effects, adjusting for age, sex, study-specific covariates, and population substructure. The threshold for genome-wide significance in the fixed-effect inverse-variance weighted meta-analysis was p<5×10^−8^. Suggestive (p<5×10^−7^) and genome-wide significant (p<5×10^−8^) variants were carried forward for replication or further investigation in an independent sample.

**RESULTS:** The discovery sample was comprised of 158,025 individuals, including 29,531 CBP cases. A genome-wide significant association was found for the intronic variant rs12310519 in *SOX5* (OR 1.08, p=7.2×10^−10^). This was subsequently replicated in an independent sample of 283,752 subjects, including 50,915 cases (OR 1.06, *p*=5.3×10^−11^), and exceeded genome-wide significance in joint meta-analysis (OR=1.07, *p*=4.5×10^−19^). We found suggestive associations at three other loci in the discovery sample, two of which exceeded genome-wide significance in joint meta-analysis: an intergenic variant, rs7833174, located between *CCDC26* and *GSDMC* (OR 1.05, p=4.4×10^−13^), and an intronic variant, rs4384683, in *DCC* (OR 0.97, p=2.4×10^−10^).

**DISCUSSION:** In this first reported meta-analysis of GWAS for CBP, we identified and replicated a genetic locus associated with CBP (*SOX5*). We also identified 2 other loci that reached genome-wide significance in a 2-stage joint meta-analysis (*CCDC26/GSDMC* and *DCC*).

## INTRODUCTION

Back pain causes more years lived with disability than any other health condition worldwide.(1) Although most adults experience a new (‘acute’) episode of back pain at some point in their lives, the societal burden of back pain is driven by the minority of individuals who fail to recover from such episodes and go on to develop persistent (‘chronic’) back pain.(2) Chronic back pain (CBP) has a number of definitions but is most often considered as back pain of duration ≥3 months in clinical practice, and a duration of ≥6 months is also commonly used in research.(3, 4)

Back pain is moderately heritable. Meta-analysis of 11 twin studies of back pain indicates a heritability of 40%, with a pattern of monozygotic (*r*_MZ_ = 0.56) and dizygotic (*r*_DZ_ = 0.28) twin correlations suggesting an additive genetic model (2*r*_DZ_ = *r*_MZ_).(5, 6) Heritability is greater for chronic than for acute back pain.(7) Nevertheless, genetic studies attempting to identify specific genetic markers for CBP have to date been limited to small studies using the candidate gene approach.(8, 9) Although CBP is often attributed to anatomic changes such as intervertebral disc degeneration or disc herniation, such findings have only weak association with CBP, (10, 11) and explain only a small proportion (7-23%) of the genetic influence on back pain(12). The unexplained genetic contribution to CBP may involve not only spine pathology but also functional predisposition to chronic pain involving higher-order neurologic processes related to the generation and maintenance of pain.(13-15) Furthermore, psychological factors such as depression are widely recognized as important risk factors for CBP.(16) Given the range of processes that might contribute to CBP, the agnostic genome-wide association approach may identify novel genetic variants associated with CBP and provide insights into underlying biological mechanisms not previously considered.

We conducted a meta-analysis of GWAS of CBP in adults of European ancestry from 16 community- and population-based cohorts. This research was an international collaboration between investigators from the Cohorts for Heart and Aging Research in Genomic Epidemiology (CHARGE) Consortium Musculoskeletal Workgroup(17) and the European Union FP7 project Pain-OMICS (‘Multi-dimensional omics approach to stratification of patients with low back pain’). The aim was to identify novel associations between specific genetic markers and CBP, and elucidate the biological mechanisms underlying this condition.

## METHODS

### Study Design and Populations

Discovery meta-analysis included adults of European ancestry from 16 population- and community-based cohorts: Cardiovascular Health Study (CHS), Framingham Heart Study (FHS), Generation Scotland (GS), Johnston County Osteoarthritis Project (JoCo), Osteoporotic Fractures in Men (MrOS) Sweden (MrOS-Gothenburg and MrOS-Malmo), MrOS US, Osteoarthritis Initiative (OAI), Rotterdam Study (RS1, RS2, and RS3), Study of Osteoporotic Fractures (SOF), 10,001 Dalmatians (Vis and Korcula), TwinsUK, and UK Biobank (UKB) participants from the interim data release(18). Replication was conducted in an independent sample of UKB European ancestry participants not included in the discovery stage, and a joint (discovery-replication) meta-analysis was performed. Detailed descriptions of the study cohorts are provided in Table 1 and the Supplemental Methods. The participating cohorts and the conduct of genetic analyses were approved by the medical ethics committees of the respective centers.

**Table 1.** Cohorts in Meta-Analysis of Genome-Wide Association Studies of Chronic Back Pain (European Ancestry)

|  |  |  |  |  |  |  |  |  |  |
| --- | --- | --- | --- | --- | --- | --- | --- | --- | --- |
| **Cohort** | **Study setting** | **Country** | **Sample size** | **Chronic back pain definition** | **Prevalence (%)** |  | **Age (yr)** | **BMI (kg/m<sup>2</sup>)** | **Women (%)** |
| Cardiovascular Health Study (CHS) | Community | USA | 2849 | ≥1 month of back pain in consecutive years | 14.2% | Cases (n=404) | 72.1 ± 5.1 | 27.3 ± 5.0 | 73.3% |
|  |  |  |  |  |  | Controls (n=2445) | 72.1 ± 5.2 | 26.1 ± 4.3 | 59.6% |
| Framingham Heart Study<sup>14</sup> | Community | USA | 2673 | ≥6 months of back pain | 21.0% | Cases (n=561) | 67.7 ± 9.3 | 28.8 ± 5.8 | 62.2% |
|  |  |  |  |  |  | Controls (n=2112) | 66.4 ± 9.1 | 28.0 ± 5.2 | 52.9% |
| Generation Scotland | Population | UK | 5071 | ≥3 months of back pain | 26.0% | Cases (n=1322) | 54.9 ± 11.8 | 28.1 ± 5.7 | 66.7% |
|  |  |  |  |  |  | Controls (n=3749) | 52.4 ± 12.7 | 26.4 ± 4.6 | 55.4% |
| Johnston County Osteoarthritis Project (JoCo) | Population | USA | 480 | ≥6 months of back pain | 38.8% | Cases (n=186) | 72.0 ± 8.0 | 31.4 ± 6.3 | 65.0% |
|  |  |  |  |  |  | Controls (n=294) | 73.0 ± 8.0 | 29.3 ± 5.2 | 58.4% |
| Mr. Os Sweden |  |  |  |  |  |  |  |  |  |
| Gothenburg | Population | Sweden | 920 | ≥6 months of back pain | 14.2% | Cases (n=131) | 75.3 ± 3.2 | 26.7 ± 3.9 | 0% |
|  |  |  |  |  |  | Controls (n=789) | 75.3 ± 3.2 | 26.1 ± 3.4 | 0% |
| Malmö | Population | Sweden | 948 | ≥6 months of back pain | 10.8% | Cases (n=102) | 75.8 ± 3.1 | 27.2 ± 3.7 | 0% |
|  |  |  |  |  |  | Controls (n=846) | 75.6 ± 3.2 | 26.4 ± 3.6 | 0% |
| Mr. Os US | Community | USA | 4615 | ≥6 months of back pain | 14.1% | Cases (n=653) | 74.6 ± 6.1 | 28.0 ± 4.1 | 0% |
|  |  |  |  |  |  | Controls (n=3962) | 73.9 ± 5.9 | 27.3 ± 3.8 | 0% |
| Osteoarthritis Initiative (OAI) | Community | USA | 2474 | ≥1 month of back pain in consecutive years | 13.5% | Cases (n=335) | 61.0 ± 9.1 | 28.9 ± 4.7 | 57.9% |
|  |  |  |  |  |  | Controls (n=2139) | 61.7 ± 9.1 | 28.1 ± 4.5 | 53.2% |
| Rotterdam Study (RS) |  |  |  |  |  |  |  |  |  |
| RS-1 | Community | Netherlands | 5965 | ≥6 months of back pain | 14.7% | Cases (n=877) | 69.1 ± 9.2 | 26.7 ± 4.0 | 72.0% |
|  |  |  |  |  |  | Controls (n=5088) | 70.0 ± 9.4 | 26.2 ± 3.7 | 58.3% |
| RS-2 | Community | Netherlands | 1566 | ≥6 months of back pain | 36.7% | Cases (n=574) | 65.3 ± 8.0 | 27.7 ± 4.2 | 66.7% |
|  |  |  |  |  |  | Controls (n=992) | 64.6 ± 8.0 | 27.3 ± 4.1 | 55.4% |
| RS-3 | Community | Netherlands | 3019 | ≥6 months of back pain | 38.2% | Cases (n=1154) | 57.4 ± 6.9 | 28.1 ± 4.8 | 59.5% |
|  |  |  |  |  |  | Controls (n=1865) | 56.9 ± 6.7 | 27.4 ± 4.5 | 54.3% |
| Study of Osteoporotic Fractures (SOF) | Community | USA | 3615 | ≥6 months of back pain | 16.3% | Cases (n=589) | 72.1 ± 5.6 | 27.6 ± 5.3 | 100% |
|  |  |  |  |  |  | Controls (n=3026) | 71.4 ± 5.2 | 26.6 ± 4.4 | 100% |
| 10,001 Dalmatians |  |  |  |  |  |  |  |  |  |
| Vis | Population | Croatia | 251 | ≥3 months of back pain | 22.3% | Cases (n=56) | 67.3 ± 13.2 | 27.8 ± 4.4 | 69.6% |
|  |  |  |  |  |  | Controls (n=195) | 63.7 ± 12.1 | 26.7 ± 4.0 | 52.8% |
| Korcula | Population | Croatia | 773 | ≥3 months of back pain | 21.2% | Cases (n=164) | 64.3 ± 12.9 | 28.0 ± 4.3 | 70.7% |
|  |  |  |  |  |  | Controls (n=609) | 58.0 ± 14.9 | 27.0 ± 4.1 | 64.0% |
| UK Biobank | Population | United Kingdom | 120,024 | ≥3 months of back pain | 18.0% | Cases (n=21,600) | 57.3 ± 7.9 | 28.5 ± 5.4 | 54.0% |
|  |  |  |  |  |  | Controls (n=98,424) | 57.0 ± 7.9 | 27.4 ± 4.8 | 52.4% |
| TwinsUK | Population- | United | 2782 | ≥3 months of back pain | 29.6% | Cases (n=823) | 56.7 ± 12.6 | 27.4 ± 5.3 | 90.3% |
|  | based twin registry | Kingdom |  |  |  | Controls (n=1959) | 54.3 ± 13.9 | 26.0 ± 4.9 | 90.0% |
| Total of all cohorts | - | - | 158,025 | - | - | Cases (n=29,531) | - | - | - |
|  |  |  |  |  |  | Controls (n=128,494) | - | - | - |

### Chronic Back Pain (CBP) Phenotype

There is no “gold-standard” definition for CBP. Consistent with the most commonly accepted clinical and research definitions for CBP (3, 4), CBP cases were defined in this study as those reporting back pain that had been present for a minimum of 3-6 months, by one of 3 definitions depending on the cohort: 1) ≥3 months of back pain, 2) ≥6 months of back pain, and 3) ≥1 month of back pain in consecutive years (see Table 1). The comparison group (“controls”) was comprised of those who reported not having back pain or reported back pain of insufficient duration to be included as a case.

### Genotyping

Details of genotyping, quality control, imputation methods, and genome-wide analysis for each cohort were study-specific (Supplemental Tables S2-3). In brief, genotyping was performed using commercially available genome-wide arrays. Imputation of single nucleotide polymorphisms (SNPs) and insertions/deletions (indels) was performed using reference panels from 1000 Genomes phase 1 version 3 or phase 3,(19) or the Haplotype Reference Consortium.(20)

### Statistical Analysis

We conducted genome-wide association analyses in each of the 16 cohorts, and subsequent meta-analysis of autosomal SNPs to combine results from all cohorts. Each site conducted GWAS using logistic regression models with additive genetic effects to test for associations between each variant and CBP as a binary trait. These models adjusted for age, sex, study-specific covariates, and population substructure using principal components (Supplemental Table S3). Height and body mass index (BMI), calculated as weight in kilograms divided by height in meters squared, were not included as covariates in site-specific GWAS, since these traits might lie along the causal pathway or (in the case of BMI) reflect a consequence of CBP. Harmonization and quality control of GWAS results from each cohort were conducted using the EasyQC software package in the R statistical environment (v3.2.2), using methods described previously.(21) After removal of SNPs with very low minor allele frequencies (MAF <0.005 to <0.03, depending on cohort) and other filters (Supplemental Methods), the number of SNPs included in the meta-analysis ranged from 6,205,227 (Croatia-Vis) to 9,775,703 (MrOS-Gothenburg) (Supplemental Table S3). Fixed-effect inverse-variance weighted meta-analysis was performed using METAL, version 2011-03-25 (http://csg.sph.umich.edu/abecasis/metal/). Quality control and meta-analysis were conducted twice, independently of each other, by researchers at the University of Washington (MP and PS) and at PolyOmica (YT, YA, and LCK). The results from the two centers were compared to ensure accuracy. We used linkage disequilibrium (LD) score regression (LDsr) to examine the potential for confounding.(22) The genome-wide significance level was defined as *p*<5×10^−8^, and suggestive significance level was defined as *p*<5×10^−7^, after using the LDsr intercept as a correction factor. Q-Q and Manhattan plots were generated in R. We conducted GCTA-COJO (Genome-Wide Complex Trait Analysis conditional and joint analyses, http://cnsgenomics.com/software/gcta/#About) to examine associations conditional on the most significant variant at each locus.

The most highly-associated variants at genome-wide significant loci were subjected to replication in the independent sample of UKB participants. Analysis in the independent sample used logistic regression with additive genetic effects, adjusting for age, sex, array, and principal components (Supplemental Table S3); significant replication was defined using a Bonferroni-corrected threshold of p<0.05 divided by the number of genome-wide significant loci. The most highly-associated variants at loci with suggestive significance were selected for a joint (discovery-replication) meta-analysis using p<5×10^−8^ to define genome-wide significance. Further details regarding analysis are provided in Supplemental Methods.

For genome-wide significant variants and those in LD (r^2^ >0.6) we examined previously reported GWAS associations with other phenoypes using PhenoScanner v1.1 (http://www.phenoscanner.medschl.cam.ac.uk/phenoscanner), and conducted functional annotation using FUMA (http://fuma.ctglab.nl). FUMA draws upon multiple publicly available databases, annotating variants for consequences on gene functions using the combined annotation dependent depletion (CADD) score,(24) potential regulatory functions (RegulomeDB score),(25) and effects on gene expression using expression quantitative trait loci (eQTLs) of different tissue types (GTExv6) (26, 27) (Supplemental File). The higher the CADD score, the more potentially deleterious is the variant. A CADD score of ≥10 indicates a variant predicted to be among the 10% most deleterious substitutions involving the human genome, a score of ≥20 indicates a variant among the 1% most deleterious, and so forth.(24) We used data from the Roadmap Epigenomics Project to evaluate whether the lead variants at each locus reside in enhancer regions for selected tissues with possible conceptual connections to the spine via roles involving chondrogenesis, vertebral development, muscle, and pain processing the central nervous system.(28, 29). Because two significant variants were found to be associated with height in prior published GWAS, we conducted *post hoc* region-specific GWAS secondary analyses accounting for height, among UKB participants from the discovery stage. Further details of functional annotation and secondary analyses are provided in Supplemental Methods.

Finally, we used LDsr of summary-level GWAS results from the discovery stage to estimate heritability due to common autosomal SNPs and genetic correlations.(22) We transformed the observed SNP heritability to the liability scale, in order to make heritability estimates for CBP comparable with traditional heritability estimates from twin studies.(30) We used cross-trait LDsr and publicly available meta-GWAS results from LDhub to examine genetic correlations with selected traits with possible links to CBP: anthropometrics (adult height, infant/childhood height, waist/hip circumference, BMI, and overweight/obesity), depression and depressive symptoms, osteoarthritis, and rheumatoid arthritis.(31, 32)

## RESULTS

### Characteristics of the discovery sample

The discovery sample included 29,531 CBP cases of the total 158,025. The characteristics of cohorts included in the meta-analysis are shown in Table 1. The mean age of participants in each cohort ranged between 53-76 years. Within cohorts, mean age, BMI, and proportion of females was more often higher among CBP cases than among those without CBP.

### Meta-analysis of GWAS of CBP

A quantile-quantile plot comparing the meta-analysis association results with those expected by chance is presented in Figure 1. The LDsr intercept was 1.007, providing no evidence of inflation of p-values from population stratification. Meta-analysis results are summarized in the Manhattan plot shown in Figure 2.

**Figure 1.**
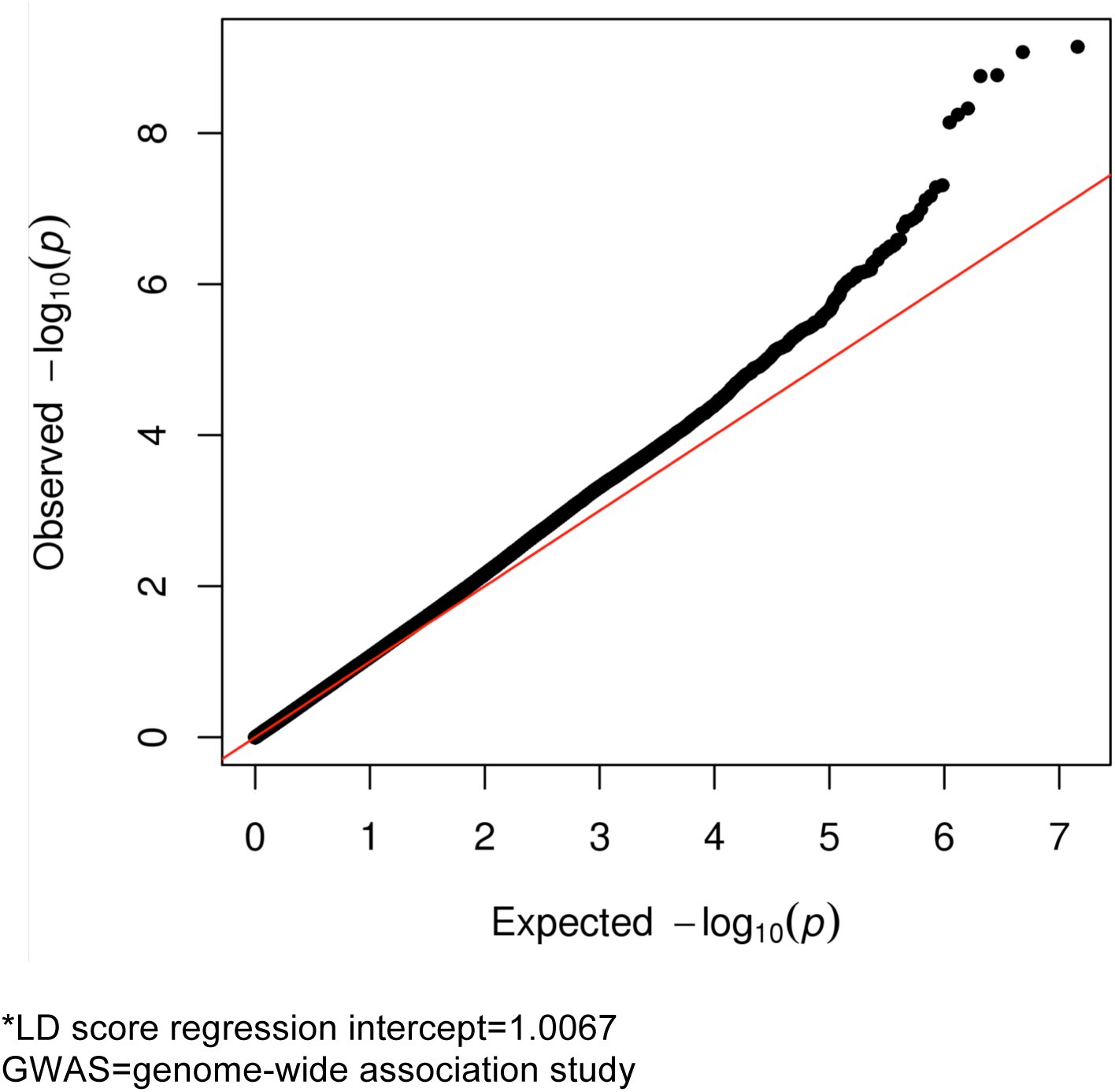
Quantile-quantile plot of p-values from the meta-analysis (discovery) of GWAS of chronic back pain (n=158,025)*

**Figure 2.**
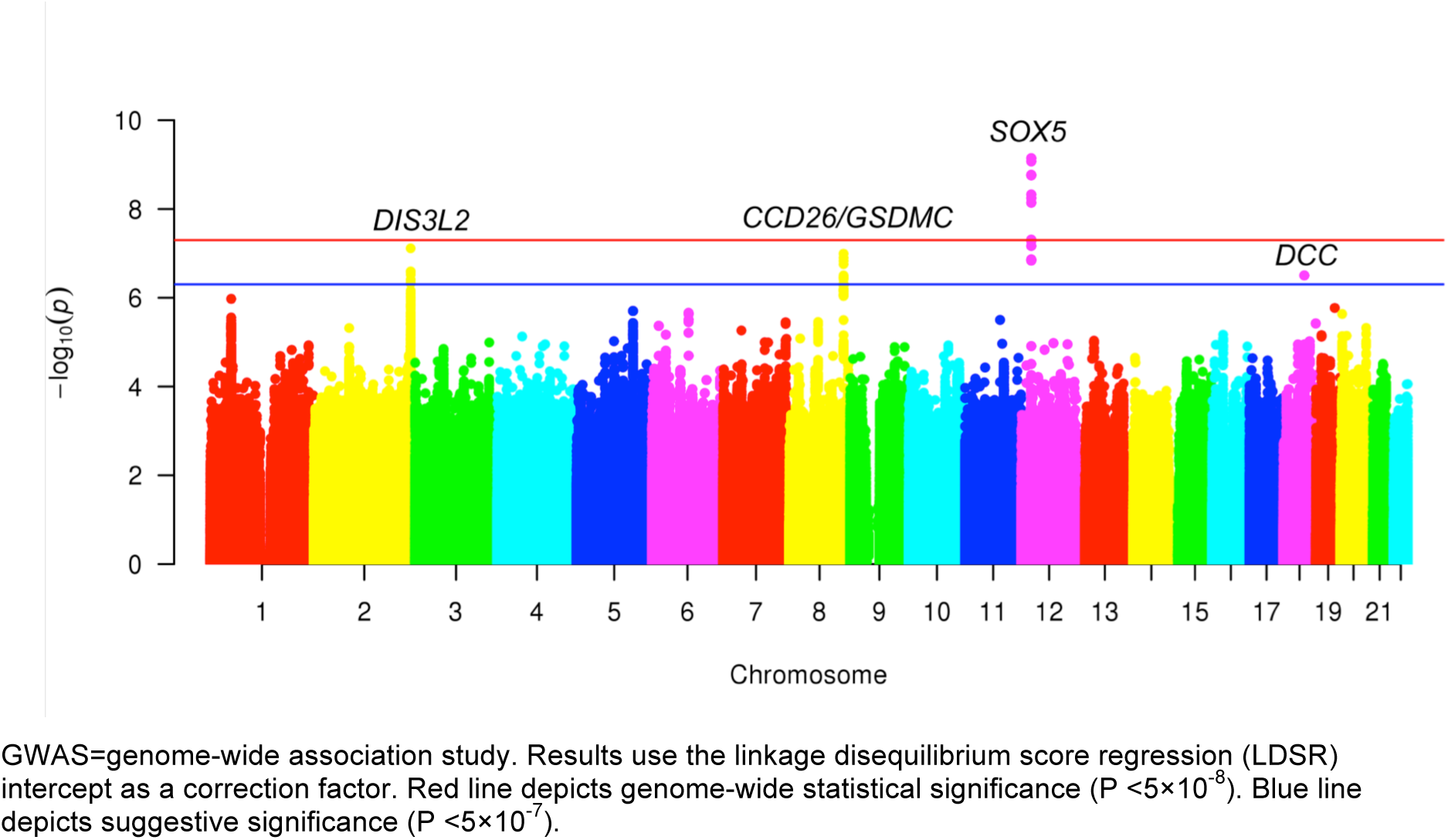
Manhattan plot for meta-analysis (discovery) of GWAS of chronic back pain (n=158,025)

A genome-wide significant association (OR 1.08, *p*=7.2×10^−10^) was found for rs12310519 on chromosome 12 in an intronic region of *SOX5*, with little evidence for heterogeneity (*I*^2^=0, *p*=0.95) (Table 2, Supplemental Figure S1). Several other signals were in high LD (*r*^2^>0.8) with the top signal (Supplemental Figure S2), but none were independently associated with CBP in analyses conditional on rs12310519.

**Table 2.** Association Results for Chronic Back Pain: Meta-analysis (Discovery), Replication, and Joint Meta-analysis*

|  |  |  |  |  |  | Meta-analysis (Discovery)<sup>a</sup> |  |  |  |  | Independent Sample (Replication)<sup>b</sup> |  | Joint Meta-Analysis (Discovery-Replication)<sup>c</sup> |  |
| --- | --- | --- | --- | --- | --- | --- | --- | --- | --- | --- | --- | --- | --- | --- |
| SNP rsID | Chr:Pos<sup>d</sup> | Nearest Gene | Location | Effect allele | Other allele | EAF | OR | p-value | I<sup>2</sup> | Het. p-value | OR | p-value<sup>b</sup> | OR | p-value |
| rs12310519<sup>e</sup> | 12:23975219 | SOX5 | intronic | T | C | 0.16 | 1.08 | 7.2 x 10<sup>-10</sup> | 0 | 0.95 | 1.06 | 5.3 x 10<sup>-11</sup> | 1.07 | 4.5 x 10<sup>-19</sup> |
| rs1453867 | 2:232917899 | DIS3L2 | intronic | T | C | 0.65 | 0.95 | 7.7 x 10<sup>-8</sup> | 13 | 0.31 | 0.98 | 0.021 | 0.97 | 3.9 x 10<sup>-7</sup> |
| rs7833174 | 8:130718772 | CCDC26/<br>GSDMC | intergenic | T | C | 0.77 | 1.06 | 1.0 x 10<sup>-7</sup> | 0 | 0.71 | 1.04 | 3.7 x 10<sup>-7</sup> | 1.05 | 4.4 x 10<sup>-13</sup> |
| rs4384683 | 18:50379032 | DCC | intronic | A | G | 0.54 | 0.95 | 3.2 x 10<sup>-7</sup> | 0 | 0.86 | 0.97 | 4.2 x 10<sup>-5</sup> | 0.97 | 2.4 x 10<sup>-10</sup> |

Chr:pos=chromosome-position, EAF=effect allele frequency OR=odds ratio, het.=heterogeneity
\*Top variant at each locus meeting suggestive or genome-wide significance level in discovery stage ( $p < 5.0 \times 10^{-7}$ )
<sup>a</sup>After genomic control using the LD score regression intercept
<sup>b</sup>Replication for rs12310519. The threshold for significance in replication of rs12310519 was $p < 0.05$ (0.05/1)
<sup>c</sup>The threshold for genome-wide significance in joint analysis was $p < 5 \times 10^{-8}$
<sup>d</sup>Build GRCh37/hg19
<sup>e</sup>rs115392701 has merged into rs12310519

No other variants achieved genome-wide significance, but variants in three other loci reached suggestive significance (Table 2, Supplemental Table S4, Supplemental Figures S3-8): rs1453867 (OR 0.95, *p*=7.7×10^−8^), located in an intronic region of chromosome 2 within *DIS3L2*; rs7833174 (OR 1.06, *p*=1.0×10^−7^), located in an intergenic region on chromosome 8 between *CCDC26* (a long non-coding RNA) and *GSDMC*; and rs4384683 (OR 0.95, *p*=3.2×10^−7^), located in an intronic region of chromosome 2 within *DCC*. In each of these 3 regions, there was no other variant reaching the suggestive significance level in analyses conditional on the lead SNP in the region.

We examined the 4 top variants in an independent sample of 283,752 UKB individuals that included 50,915 cases (Table 2). For all 4 variants, the direction of association was the same in discovery and replication. The association for rs12310519 in *SOX5* replicated in the independent sample (OR 1.06, *p*=5.3×10^−11^), and exceeded genome-wide significance in the joint analysis (OR=1.07, *p*=4.5×10^−19^). Of the 3 suggestive-significance variants from the discovery stage, rs7833174 at *CCDC26/GSDMC* (OR 1.05, *p*=4.4×10^−13^) and rs4384683 in *DCC* (OR 0.97, *p*=2.4×10^−10^) exceeded genome-wide significance in the joint meta-analysis, but rs1453867 in *DISL32* (OR 0.98, *p*=3.9×10^−7^) did not (Table 2). Thus, we demonstrate genome-wide significant associations of CBP with loci tagged by rs12310519 (*SOX5),* rs7833174 (*CCDC26/GSDMC*), and rs4384683 (*DCC*), with replication for rs12310519 in *SOX5*.

### Characterization of variants in SOX5, CCDC26/GSDMC, and DCC

The variant rs9804988, in LD with the lead SNP in *SOX5* rs12310519, was associated with height (*p*=9.6×10^−6^) in a prior GWAS (Supplemental Table S5). The T allele at rs9804988 was associated both with greater height and greater CBP risk (OR = 1.08). The highest CADD score among *SOX5* variants was 10.52 for the lead SNP in the region rs12310519, indicating it is predicted to be among the 10% most deleterious possible substitutions in the human genome (Supplemental File). However, the overall regulatory potential of these variants was low according to RegulomeDB score (scores of 6 [‘minimal binding evidence’]). The lead SNP rs12310519 and variants in LD contain active enhancer marks in chondrogenic cells (Figure 3).

**Figure 3.**
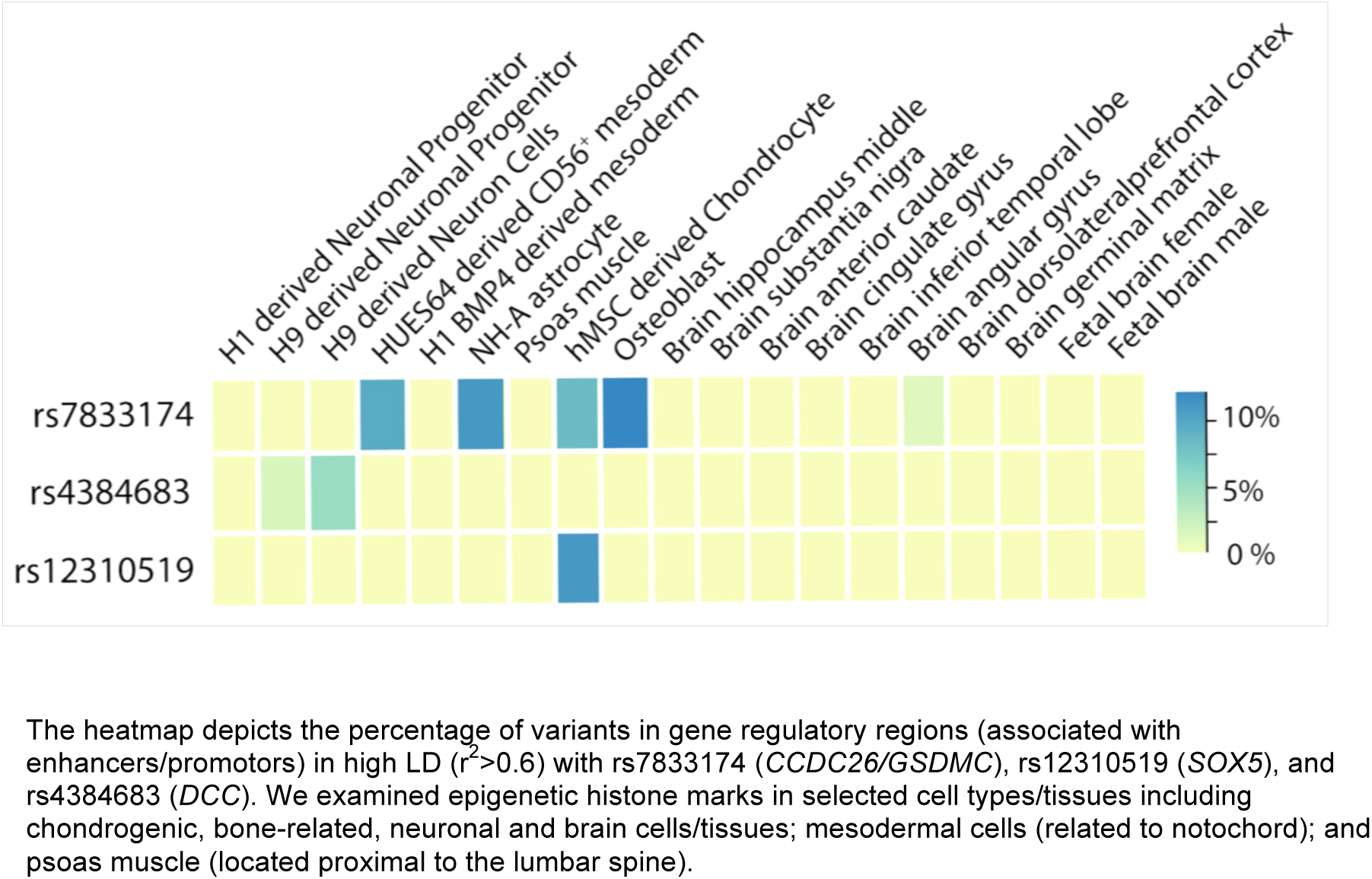
Significant variants are co-localized with potential gene regulatory markers

Variants in the intergenic *CCDC26/GSDMC* locus were significantly associated with height in prior GWAS (lowest *p*=1.0×10^−41^, Supplemental Table S5); alleles associated with greater height were also associated with greater CBP risk. Variants in *CCDC26/GSDMC* that were suggestively associated with CBP were also associated with lumbar microdiscectomy for sciatica in a recent GWAS of Icelandic adults (Supplemental Table S6, lowest *p*=5.6×10^−12^), with the same direction of effect (i.e. alleles associated with greater risk of lumbar discectomy for sciatica were also associated with greater CBP risk). The highest CADD score among these variants was 18.75 for rs6470778, indicating that this SNP is predicted to be among the 5% most deleterious substitutions in the human genome, and the overall regulatory potential of these variants was substantial according to Regulome DB score (highest RegulomeDB score of 2b [‘likely to affect binding’]) (Supplemental File). These variants were also eQTLs for *GSDMC* expression in esophageal mucosa, skin, and skeletal muscle (Supplemental File, *p*<5×10^−8^). The lead SNP rs7833174 and variants in LD contain active enhancer marks located in regulatory regions for mesodermal cells, astrocytes, chondrogenic cells, and osteoblasts (Figure 3).

No variants in the intronic *DCC* locus were associated with other phenotypes in prior GWAS. The highest CADD score among these variants was 11.21 for rs2116378, indicating that it is predicted to be among the 10% most deleterious substitutions in the human genome, but the overall regulatory potential of these variants was low according to Regulome DB score (Supplemental File, highest RegulomeDB score of 5). The lead SNP rs4384683 and variants in LD contain active enhancer marks in H9 human embryonic stem cell-derived neural cells (Figure 3).

### Secondary analyses to examine interrelationships with height

*Post hoc* analyses among UKB participants from the discovery sample (n=120,023) indicated that associations with CBP for the lead variant in *SOX5* were similar with and without adjustment for height as a covariate, and in conditional analyses accounting for height (Supplemental Table S7). Associations with CBP for the lead variant in *CCDC26/GSDMC* were also similar with and without adjustment for height as a covariate. However, associations with CBP were markedly diminished when conditional on the lead height-associated variant in the region, and associations with height were markedly diminished when conditioned on the lead CBP-associated variant in the region. This suggests that the same functional variant is responsible for association of *CCDC26/GSDMC* locus with height and CBP, although an alternative explanation is two functional variants in tight LD.

### Heritability of CBP and genetic correlations

SNP heritability of CBP on the liability scale was 7.6%. Genetic correlations of nominal significance (range of *r*_g_ 0.17- 0.31, p<0.05) were found with anthropometric traits involving obesity or body fat distribution (waist circumference, hip circumference, waist-hip ratios, overweight/obesity classes, and BMI), but not with height (Supplemental Table S8). Larger magnitude nominally significant (p<0.05) genetic correlations were also found with depression-related phenotypes (range of *r*_g_ 0.46-0.52), self-reported osteoarthritis (*r*_g_=0.63), and ICD-10-defined osteoarthritis (*r*_g_=0.49) phenotypes.

## DISCUSSION

This study is the first meta-GWAS of CBP. This collaboration between two international consortia for genomic studies of complex traits in the USA and Europe incorporated data from 16 cohorts and more than 441,000 participants of European ancestry across discovery and replication samples. Our study identifies three novel associations with CBP for loci at *SOX5, CCDC26/GSDMC,* and *DCC*.

CBP was most strongly associated with rs12310519 in an intronic region of the *SOX5* gene. The *SOX* genes are a family of transcription factors involved in virtually all phases of embryonic development, and are thought to determine the fate of many cell types. The *SOX* genes are defined by containing the HMG (‘high mobility group’) box of a gene involved in sex determination called SRY (‘sex determining region’) (33). *SOX5* and *SOX6* have overlapping functions and work together in close coordination that is necessary for efficient chondrogenesis (34). Inactivation of *SOX5* leads to minor defects in cartilage and skeletogenesis in mice, whereas *SOX5/SOX6* double knockouts have severe chondrodysplasia (35). Together with *SOX9*, *SOX5* and *SOX6* are sometimes referred to as the ‘master chondrogenic SOX trio’ (34, 36). Prior work indicates an important role for *SOX5* in articular cartilage and osteoarthritis (37, 38), and such a role was also supported by our functional annotation showing that rs12310519 (and SNPs in high LD) overlapped with potential regulatory regions for chondrogenic cells. *SOX5/6* are also essential for notochord development, and through this role they are critical in the formation of the vertebral column, including the intervertebral discs (39). Inactivation of *SOX5* and/or *SOX6* in mice leads to a range of abnormalities in the development of spinal structures (39). Although variants in *SOX5* have not been reported in prior GWAS of limb osteoarthritis (knee, hip, or hand) (40-47), the association of *SOX5* with CBP may involve the spinal structures specifically. *SOX5* was also not associated with lumbar intervertebral disc degeneration in the prior two GWAS conducted for intervertebral disc degeneration (48, 49). However, further GWAS of disc-related phenotypes are much needed, as are GWAS of the zygapophyseal (‘facet’) joint osteoarthritis, the only synovial joints in the spine (50).

The intergenic variants at *CCDC26/GSDMC* associated with CBP in the current study were also associated with lumbar microdiscectomy for sciatica in a prior GWAS, with the same direction of effect.(51) These findings are intriguing, given that lumbar disc herniations have long been implicated as a cause of some forms of back pain.(52) Recent studies have concluded that associations between imaging-detected lumbar disc herniation and CBP are of modest magnitude.(10, 11) This might explain the small magnitude association of the top variant at *CCDC26/GSDMC* with CBP in the current study (OR 1.08 in discovery), in contrast to the larger magnitude association seen with microdiscectomy of sciatica (OR 1.23). Functional characterization of these intergenic variants indicates the likely involvement of the gene *GSDMC*. *GSDMC* encodes the protein Gasdermin C, part of the *GSDM* family of genes that is expressed in epithelial tissues. Although the specific role of *GSDMC* in lumbar disc herniation and/or sciatica is unclear, *GSDMC* is associated with differential methylation patterns in osteoarthritis-related cartilage and subchondral bone cartilage, (53, 54) consistent with our finding that variants in LD with rs4384683 were located in potential regulatory regions in chondrocytes and osteoblasts. Taken together, these data suggest interconnections between variants at *CCDC26/GSDMC* and CBP involving cartilage, osteoarthritis, and/or intervertebral disc degeneration.

The third significant CBP-associated variant in our study was rs2116378, an intronic variant in the gene *DCC* (Deleted in Colorectal Carcinoma), which co-localized with regulatory regions in neural embryonic stem cells. *DCC* encodes a transmembrane protein that is a receptor for netrin-1, an axonal guidance molecule involved in the development of spinal and cortical commissural neurons.(55) Interactions between DCC and netrin-1 are among the best-studied axonal guidance processes, with key roles during development and in adulthood, and they also affect angiogenesis.(56, 57) Increased expression of netrin-1 and DCC occurs in degenerate human intervertebral discs compared to healthy control discs, and in nucleus pulposus compared to annulus fibrosis.(58) Netrin-1/DCC might therefore mediate neurovascular ingrowth into the intervertebral disc, which has long been implicated as a possible mechanism of chronic discogenic back pain.(58, 59) Netrin-1/DCC interactions also play a role in pain processing in the spinal cord in animal models of mechanical allodynia.(56) Taken together, these findings suggest nociceptive and/or neuropathic mechanisms connecting lumbar intervertebral disc degeneration (including disc herniation) with CBP.

Variants in *CCDC26/GSDMC* and *SOX5* associated with greater risk of CBP in our meta-analysis were also reported to be associated with greater height in prior GWAS. Our secondary analyses showed that the association of *SOX5* variants with CBP was independent of height, however, CBP- and height-associated variants in *CCDC26/GSDMC* were tightly linked and could not be disentangled in conditional analyses. The association of variants in *CCDC26/GSDMC* with both CBP and height might be explained by pleiotropy, causal connections, or other factors. The direction of effects in our study for *CCDC26/GSDMC* variants were consistent with epidemiological reports that greater height confers increased risk of back pain (60-62), although a systematic review found no such association.(63) These findings might be viewed as supporting a causal link between height and CBP, but could instead simply reflect shared underlying genetic factors between the two phenotypes. CBP and height have compelling connections to the spine. Prior studies demonstrating the vital role of *SOX5* in normal vertebrate development (34, 36, 39) may hold lessons that apply to other genetic markers including those in *CCDC26/GSDMC*: if associations with CBP and height are mediated by the spine and/or vertebral column development, it will be difficult to distinguish pleiotropy and causality using GWAS alone.

SNP-based heritability in the current study (8%) was considerably lower than estimates from twin studies (~40%). This is a common situation with modern methods of estimating heritability using genotype data, since such estimates reflect only one aspect of narrow-sense heritability captured by the additive genetic components of common variants, excluding the contributions of rare variants, non-additive effects, epistasis, or gene-environment interactions.(30) Despite the modest heritability of CBP (and other self-reported traits), we found significant and large magnitude genetic correlations between CBP and other phenotypes that may be risk factors for CBP or consequences of CBP, such as depression and obesity-related factors (but not height). Future GWAS of CBP may benefit from taking these relationships into account, either by accounting for these phenotypes as covariates, or in multivariate GWAS designs.

A distinguishing feature of the current study as compared to many other GWAS is that the CBP phenotype examined represents a symptom, rather than a disease or a biomarker. Although successful GWAS of self-reported symptoms have been conducted which replicate associations seen with more specific disease phenotypes,(64) our findings highlight potential challenges of GWAS of CBP: despite being one of the largest international studies of CBP ever conducted, our study detected only 3 significant associations with CBP. Still larger sample sizes may be needed in future discovery efforts using this phenotype, or different methods should be used. A consequence of the nonspecific nature of the CBP phenotype is the fact that, unlike other musculoskeletal phenotypes such as osteoarthritis, the tissue correlates optimal for conducting functional follow-up studies of findings from CBP GWAS are unclear. Most animal models for back pain rely on specific mechanisms of inducing pain, such as injuries to the intervertebral disc, zygapophyseal (‘facet’) joint, dorsal root ganglion, or muscle.(65) However, each of these mechanisms likely explain only a certain portion of back pain cases, and do not encompass the important psychosocial aspects of pain and pain reporting that are relevant in humans. Despite the importance of psychosocial factors, our meta-GWAS findings are a reminder that structural/anatomic factors involving spinal degeneration, such as disc herniations or osteoarthritis of spinal structures (e.g. facet joints), remain potentially important contributors to the CBP. Future GWAS of CBP may benefit from alternative strategies such as CBP endophenotyping (stratification by back pain subtypes), simultaneously studying CBP and spinal degeneration/fracture phenotypes, and allowing for the examination of interactions between genetic markers for spinal degeneration and markers for pain processing or axonal signaling (including *DCC* and netrin-1).

Strengths of our study include its multicohort design and large sample size. A potential limitation of our study was heterogeneity of the CBP phenotype used, a consequence of pooling data from numerous cohorts using different definitions. Although this approach may have helped identify genetic associations shared across CBP subtypes, it might obscure associations pertinent to specific subtypes of back pain. Despite phenotype heterogeneity, which would be expected to bias towards the null, our study successfully identified several associations of statistical significance. Recent efforts to standardize CBP definitions may help to limit phenotype heterogeneity in future meta-GWAS of CBP.(3) Another aspect of the phenotype used in our meta-analysis was that individuals with back pain of less than 3-6 months duration were included as controls. This was done deliberately so as to focus on back pain of chronic duration as the phenotype of interest. That said, GWAS examining back pain of *any* duration, or analyses excluding those with non-chronic back pain, may find different results from ours.

In summary, this meta-analysis of GWAS of CBP identified novel genetic associations with CBP at *SOX5*, *CCDC26/GSDMC,* and *DCC.* These findings suggest possible mechanisms involving the spine and/or shared biological pathways involving CBP, cartilage, lumbar disc herniation, height and/or vertebral development.

## ACKNOWLEDGEMENTS

Infrastructure for the CHARGE Consortium is supported in part by the National Heart, Lung, and Blood Institute grants R01HL105756.

This study was supported by the European Commissions’ Seventh Framework Programme funded project PainOmics (Grant agreement n. 602736)

**Cardiovascular Health Study:** This CHS research was supported by NHLBI contracts HHSN268201200036C, HHSN268200800007C, HHSN268200960009C, N01HC55222, N01HC85079, N01HC85080, N01HC85081, N01HC85082, N01HC85083, N01HC85086; and NHLBI grants U01HL080295, R01HL087652, R01HL105756, R01HL103612, R01HL120393, R01HL085251, and R01HL130114 with additional contribution from the National Institute of Neurological Disorders and Stroke (NINDS). Additional support was provided through R01AG023629 from the National Institute on Aging (NIA). A full list of principal CHS investigators and institutions can be found at <u>CHS-NHLBI.org</u>. The provision of genotyping data was supported in part by the National Center for Advancing Translational Sciences, CTSI grant UL1TR000124, and the National Institute of Diabetes and Digestive and Kidney Disease Diabetes Research Center (DRC) grant DK063491 to the Southern California Diabetes Endocrinology Research Center. The content is solely the responsibility of the authors and does not necessarily represent the official views of the National Institutes of Health.

**Framingham Heart Study:** From the Framingham Heart Study of the National Heart Lung and Blood Institute of the National Institutes of Health and Boston University School of Medicine. This work was supported by the National Heart, Lung and Blood Institute's Framingham Heart Study (Contract No. N01-HC-25195) and its contract with Affymetrix, Inc for genotyping services (Contract No. N02-HL-6-4278), and by grants from the National Institute of Neurological Disorders and Stroke (NS17950) and the National Institute of Aging, (AG08122, AG16495). The content is solely the responsibility of the authors and does not necessarily represent the official views of the Framingham Heart Study or Boston University School of Medicine.

**Generation Scotland:** Generation Scotland (GS) received core funding from the Chief Scientist Office of the Scottish Government Health Directorates CZD/16/6 and the Scottish Funding Council HR03006. Genotyping of samples was carried out by the Genetics Core Laboratory at the Wellcome Trust Clinical Research Facility, Edinburgh, Scotland and was funded by the Medical Research Council UK and the Wellcome Trust (Wellcome Trust Strategic Award “STratifying Resilience and Depression Longitudinally” (STRADL) Reference 104036/Z/14/Z).

**Johnston County Osteoarthritis Project:** The JoCo is supported in part by S043, S1734, & S3486 from the CDC/Association of Schools of Public Health; 5-P60-AR30701 & 5-P60-AR49465-03 from NIAMS/NIH; genotyping was supported by Algynomics, Inc.

**Mr. Os Sweden:** MrOS Sweden is supported by the Swedish Research Council, the Swedish Foundation for Strategic Research, the ALF/LUA research grant in Gothenburg, the Lundberg Foundation, the Knut and Alice Wallenberg Foundation, the Torsten Soderberg Foundation, and the Novo Nordisk Foundation, the ALF/FoUU research grant in Skane, Herman Järnhardts, Kocks and Österluds Foundations.

**Osteoporotic Fractures in Men (Mr Os) US:** The Osteoporotic Fractures in Men (MrOS) Study is supported by National Institutes of Health funding. The following institutes provide support: the National Institute of Arthritis and Musculoskeletal and Skin Diseases (NIAMS), the National Institute on Aging (NIA), the National Center for Research Resources (NCRR), and NIH Roadmap for Medical Research under the following grant numbers: U01 AR45580, U01 AR45614, U01 AR45632, U01 AR45647, U01 AR45654, U01 AR45583, U01 AG18197, U01-AG027810, U01 AG042140, U01 AG042143, U01 AG042124, U01 AG042145, U01 AG042139, U01 AG042168, U01 AR066160, UL1 TR000128, and UL1 RR024140. The National Institute of Arthritis and Musculoskeletal and Skin Diseases (NIAMS) provides funding for the MrOS ancillary study ‘GWAS in MrOS and SOF’ under the grant number RC2ARO58973.

**Osteoarthritis Initiative (OAI):** This study was supported by the American Recovery and Reinvestment Act (ARRA) through grant number RC2-AR-058950 from NIAMS/NIH. The OAI is public-private partnership comprised of five contracts (N01-AR-2-2258; N01-AR-2-2259; N01-AR-2-2260; N01-AR-2-2261; N01-AR-2-2262) funded by the NIH. Additional support was provided by NIH grant P30-DK072488.

**Rotterdam Study:** The Rotterdam Study is supported by the Erasmus Medical Center and Erasmus University, Rotterdam; the Netherlands Organization for Scientific Research (NWO), the Netherlands Organization for Health Research and Development (ZonMw), the Research Institute for Diseases in the Elderly (RIDE), the Ministry of Education, Culture and Science, the Ministry for Health, Welfare and Sports, the European Commission (DG XII), and the Municipality of Rotterdam. The generation and management of GWAS genotype data for the Rotterdam Study (RS I, RS II, RS III) was executed by the Human Genotyping Facility of the Genetic Laboratory of the Department of Internal Medicine, Erasmus MC, Rotterdam, The Netherlands. The GWAS datasets are supported by the Netherlands Organisation of Scientific Research NWO Investments (nr. 175.010.2005.011, 911-03-012), the Genetic Laboratory of the Department of Internal Medicine, Erasmus MC, the Research Institute for Diseases in the Elderly (014-93-015; RIDE2), the Netherlands Genomics Initiative (NGI)/Netherlands Organisation for Scientific Research (NWO) Netherlands Consortium for Healthy Aging (NCHA), project nr. 050-060-810. We thank Pascal Arp, Mila Jhamai, Marijn Verkerk, Lizbeth Herrera and Marjolein Peters, MSc, and Carolina Medina-Gomez, MSc, for their help in creating the GWAS database, and Karol Estrada, PhD, and Carolina Medina-Gomez, MSc, for the creation and analysis of imputed data

**Study of Osteoporotic Fractures:** The Study of Osteoporotic Fractures (SOF) is supported by National Institutes of Health funding. The National Institute on Aging (NIA) provides support under the following grant numbers: R01 AG005407, R01 AR35582, R01 AR35583, R01 AR35584, R01 AG005394, R01 AG027574, and R01 AG027576. The National Institute of Arthritis and Musculoskeletal and Skin Diseases (NIAMS) provides funding for the MrOS ancillary study ‘GWAS in MrOS and SOF’ under the grant number RC2ARO58973.

**10,001 Dalmatians:** We would like to acknowledge the staff of several institutions in Croatia that supported the field work, including but not limited to The University of Split and Zagreb Medical Schools, the Institute for Antropological Research in Zagreb and the Croatian Institute for Public Health. We would like to acknowledge the invaluable contributions of the recruitment team in Korcula, the administrative teams in Croatia and Edinburgh and the participants. The SNP genotyping for the Vis cohort was performed in the core genotyping laboratory of the Wellcome Trust Clinical Research Facility at the Western General Hospital, Edinburgh, Scotland. The SNP genotyping for the Korcula cohort was performed in Helmholtz Zentrum München, Neuherberg, Germany. The SNP genotyping was performed by AROS Applied Biotechnology, Aarhus, Denmark. The study was funded by the Medical Research Council UK, The Croatian Ministry of Science, Education and Sports (grant 216-1080315-0302), the European Union framework program 6 EUROSPAN project (contract no. LSHG-CT-2006-018947), the Croatian Science Foundation (grant 8875), the Centre for Research Excellence in Personalized Medicine and the Centre of Competencies for Integrative Treatment, Prevention and Rehabilitation using TMS.

**TwinsUK:** TwinsUK receives funding from the Wellcome Trust; European Community’s Seventh Framework Programme (FP7/2007-2013 to Twins UK); the National Institute for Health Research (NIHR) Clinical Research Facility at Guy’s & St Thomas’ NHS Foundation Trust and NIHR Biomedical Research Centre based at Guy’s and St Thomas’ NHS Foundation Trust and King’s College London.

**UK Biobank:** This research has been conducted using the UK Biobank Resource (project # 18219). We are grateful to the UK Biobank participants for making such research possible.

Dr. Suri’s time for this work was supported by VA Career Development Award # 1IK2RX001515 from the United States (U.S.) Department of Veterans Affairs Rehabilitation Research and Development Service. Dr. Suri is a Staff Physician at the VA Puget Sound Health Care System. Dr. Smith is a Research Scientist at the VA Puget Sound Health Care System. The contents of this work do not represent the views of the U.S. Department of Veterans Affairs or the United States Government. The work of Dr. Tsepilov was supported by was supported by the Russian Ministry of Science and Education under the 5-100 Excellence Programme.

---

**Supplemental Table S1.** Chronic back pain definitions and related question items

|  |  |  |
| --- | --- | --- |
| **Cohort** | **Chronic back pain definition** | **Back pain questions for each cohort** |
| Cardiovascular Health Study | ≥1 month of back pain in consecutive years | ***“In the past year, have you had pain in your back for more than half the days of any month?”***<br>“Yes” in ≥2 consecutive years (**cases**) vs. other combinations of non-missing responses in consecutive years (**controls**) |
| Framingham Heart Study | ≥6 months of back pain | ***“Have you had back pain in the past 12 months?”***<br>“Most of the days”/“all days” (**cases**) vs. “no”/“a few days”/“some of the days” (**controls**) |
| Generation Scotland | ≥3 months of back pain | Question combinations indicating “***Have you been troubled by pain or discomfort*** [in the back], ***either all the time or on and off...for more than 3 months?***”<br>“Yes” (**cases**) vs. “No” (**controls**) |
| Johnston County Osteoarthritis Project | ≥6 months of back pain | ***“On MOST days of ANY ONE MONTH in the LAST 12 MONTHS did you have pain, aching, or stiffness in any of the following?”***<br>“Lower back (L1 through S1)” and “middle back (thoracic)” locations were specified.<br>≥6 month duration “with pain on most days” (**cases**) vs. other non-missing response options (**controls**) |
| Mr. Os Sweden |  |  |
| Gothenburg | ≥6 months of back pain | ***“Have you had back pain in the past 12 months?”***<br>“Most of the time”/“all of the time” (**cases**) vs. “no”/“never”/“rarely”/“some of the time” (**controls**) |
| Malmö | ≥6 months of back pain | ***“Have you had back pain in the past 12 months?”***<br>“Most of the time”/“all of the time” (**cases**) vs. “no”/“never”/“rarely”/“some of the time” (**controls**) |
| Mr. Os US | ≥6 months of back pain | ***“During the past 12 months have you experienced any back pain?”***<br>“Most of the time”/“all of the time” (**cases**) vs. “no”/“never”/“rarely”/“some of the time” (**controls**) |
| Osteoarthritis Initiative | ≥1 month of back pain in consecutive years | ***“How often were you bothered by back pain in the past 30 days?”***<br>“All of the time”/“Most of the time” in ≥2 consecutive years (**cases**) vs. any other combination of non-missing responses in consecutive years (**controls**) |
| Rotterdam Study (RS) |  |  |
| RS-1 | ≥6 months of back pain | ***“Have you had pain in the low back in the last month?”***<br>≥6 month duration of low back pain (**cases**) vs. no back pain or <6 months duration low back pain (**controls**) |
| RS-2 | ≥6 months of back pain | ***“Do you have pain or stiffness in your back?”***<br>≥6 month duration (**cases**) vs. no back pain/stiffness or <6 months duration back pain/stiffness (**controls**) |
| RS-3 | ≥6 months of back pain | ***“Do you have pain or stiffness in your back?”***<br>≥6 month duration (**cases**) vs. no back pain/stiffness or <6 months duration back pain/stiffness (**controls**) |
| Study of Osteoporotic Fractures | ≥6 months of back pain | ***“During the past 12 months have you experienced any back pain?”***<br>“Most of the time”/“all of the time” (**cases**) vs. “no”/“never”/“rarely”/“some of the time” (**controls**) |
| 10,001 Dalmatians |  |  |
| Vis | ≥3 months of back pain | ***“Have you ever had an episode of chronic LBP that lasted for over 3 months?”*** and ***“Do you have low back pain at the moment?”***<br>“Yes” to both questions (**cases**) vs. “No” to either question (**controls**) |
| Korcula | ≥3 months of back pain | ***“Have you ever had an episode of chronic LBP that lasted for over 3 months?”*** and ***“Do you have low back pain at the moment?”***<br>“Yes” to both questions (**cases**) vs. “No” to either question (**controls**) |
| TwinsUK | ≥3 months of back pain | ***“In the past 3 months, have you had pain in your back on most days?”*** or ***“Have you had pain...in this area*** [lumbar or thoracic] ***for at least the past 3 months?”***<br>“Yes” to either question (**cases**) vs. “No” to both questions (**controls**) |
| UK Biobank | ≥3 months of back pain | ***“In the last month have you experienced any of the following that interfered with your usual activities?”*** (Back pain is a response option)<br>***“Have you had back pain for more than 3 months?”***<br>“Yes” to both questions (**cases**) vs. “No” to the first question or “Yes” to the first question and “No” to the second question (**controls**) |

**Supplemental Table S2.**
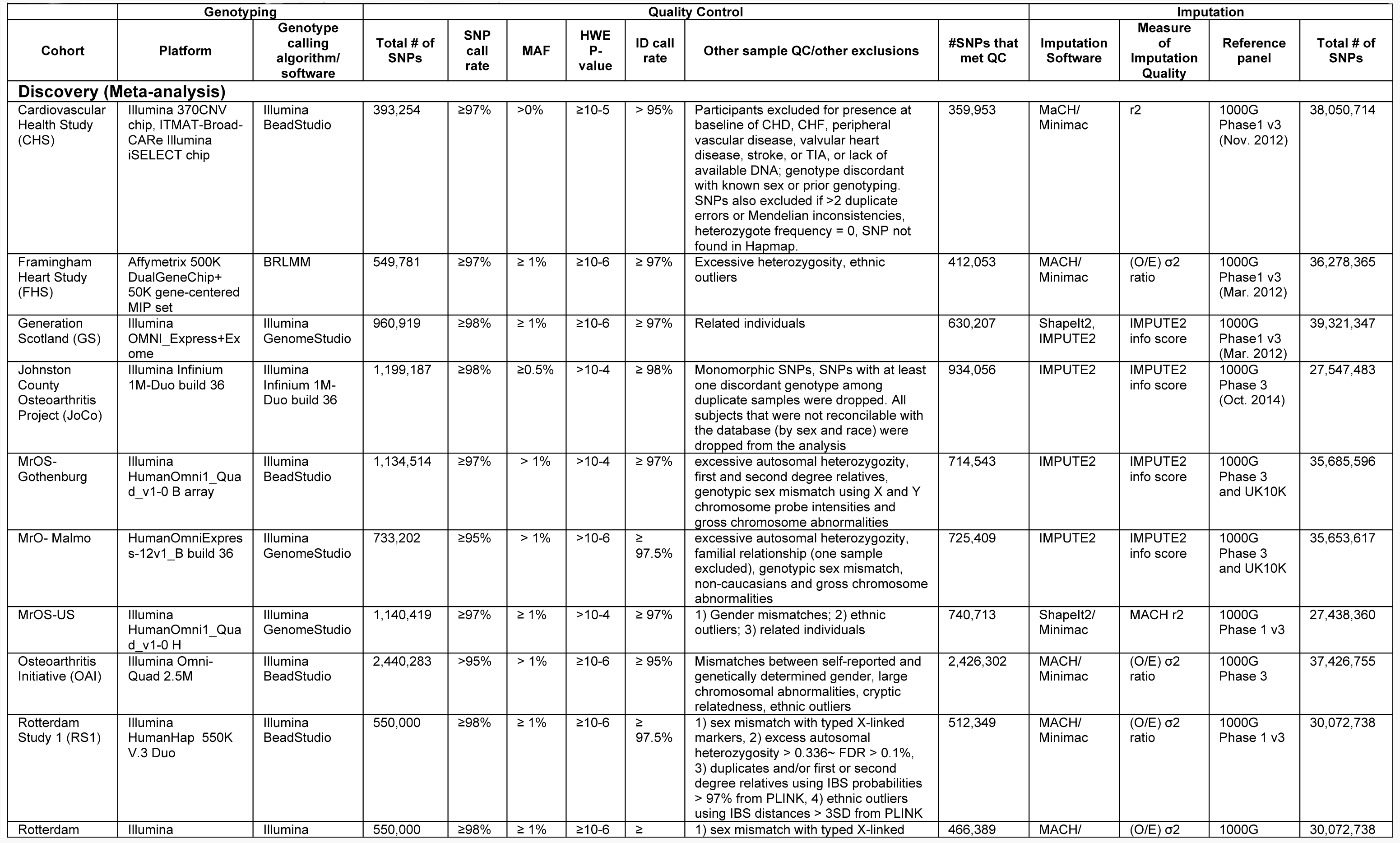
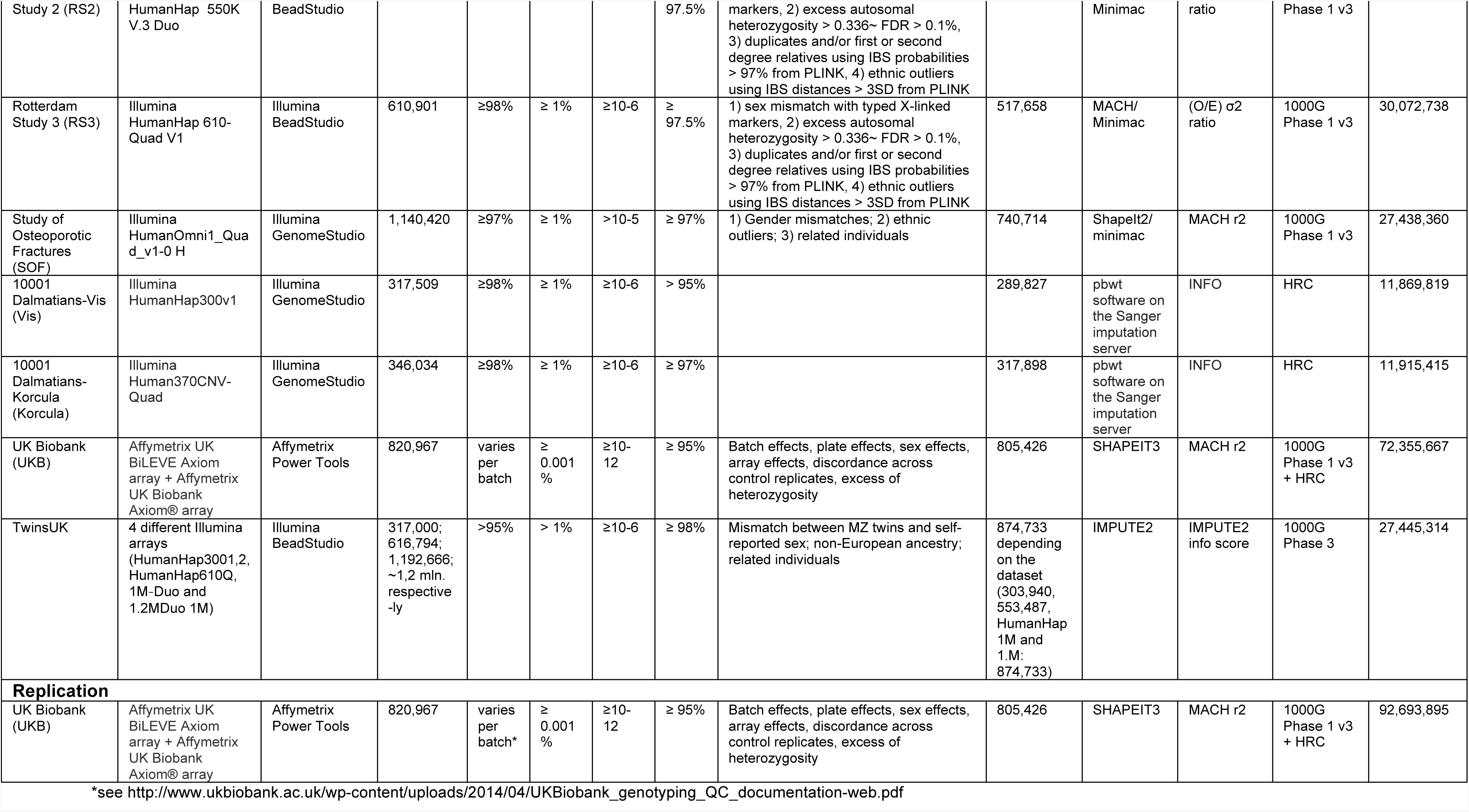
Genotyping, quality control, and imputation for each cohort

**Supplemental Table S3.** Details of the GWAS analysis for each cohort and quality control prior to meta-analysis

|  |  |  |  |  |  |  |  |  |  |
| --- | --- | --- | --- | --- | --- | --- | --- | --- | --- |
| Cohort | Total # of SNPs post-imputation | Filters |  | Model used | # SNPs in analysis | Method used to correct for genetic substructure | Software | Following file-level and meta-level quality control (see Supplemental Methods)<sup>a</sup> |  |
| | | MAF | Imputation quality | | | | | # SNPs in meta-analysis | Genomic Inflation Factor ($\lambda_{GC}$) |
| Discovery (Meta-analysis)<sup>a</sup> |  |  |  |  |  |  |  |  |  |
| Cardiovascular Health Study (CHS) | 38,050,714 | > 0% | variance on allele dosage ≤0.01 | SNP+ age + sex + clinic | 9,854,835 | NA | R | 7,528,846 | 1.00 |
| Framingham Heart Study (FHS) | 36,278,365 | > 0% | ≥0.3 | SNP + age + sex + PC1 + PC2 + PC3 + PC4 + PC5 + kinship | 19,344,131 | EIGENSTRAT | R/GEE | 7,114,199 | 1.03 |
| Generation Scotland (GS) | 39,321,347 | > 0% | none | SNP + age + sex (Unrelated individuals only) | 39,321,347 | Unrelated only | R/ProbABEL | 8,934,663 | 1.01 |
| Johnston County Osteoarthritis Project (JoCo) | 27,547,483 | > 0% | none | SNP + age + sex + PC1 + PC2 + PC3 | 27,547,483 | EIGENSOFT | PLINK | 9,236,276 | 1.02 |
| MrOS-Gothenburg | 35,685,596 | > 0% | >0.3 | SNP + age | 18,648,740 | NA | PLINK | 9,775,703 | 1.00 |
| MrO- Malmö | 35,653,617 | > 0% | >0.3 | SNP + age | 19,943,738 | NA | PLINK | 9,764,582 | 1.00 |
| MrOS-US | 27,438,360 | none | none | SNP + age + clinic site + PC1 + PC2 + PC3 + PC4 | 27,438,360 | R/GWASTools package | PLINK | 8,579,487 | 1.00 |
| Osteoarthritis Initiative (OAI) | 37,426,755 | > 0% | >0.3 | SNP + age + sex + PC1 + PC2 | 8,515,357 | PLINK-MDS | PLINK | 7,552,530 | 1.01 |
| Rotterdam Study 1 (RS1) | 30,072,738 | > 0% | none | SNP+ age + sex | 30,072,738 | NA | MACH/GRIMP | 8,017,540 | 1.01 |
| Rotterdam Study 2 (RS2) | 30,072,738 | > 0% | none | SNP+ age + sex | 30,072,738 | NA | MACH/GRIMP | 7,987,628 | 1.00 |
| Rotterdam Study 3 (RS3) | 30,072,738 | > 0% | none | SNP+ age + sex | 30,072,738 | NA | MACH/GRIMP | 8,055,340 | 1.02 |
| Study of Osteoporotic Fractures (SOF) | 27,438,360 | none | none | SNP + age+ clinic site + PC1 + PC2 + PC3 + PC4 | 27,438,360 | R/GWASTools package | PLINK | 8,570,668 | 1.02 |
| 10001 Dalmatians-Korcula (Korcula) | 11,915,415 | > 0% | none | SNP + age + sex + kinship (PC1 + PC2) | 11,915,415 | GenABEL kinship matrix | RegScan+SNPTEST | 7,814,464 | 1.01 |
| 10001 Dalmatians-Vis (Vis) | 11,869,819 | > 0% | none | SNP + age + sex + kinship (PC1 + PC2) | 11,869,819 | GenABEL kinship matrix | RegScan+SNPTEST | 6,205,227 | 1.01 |
| UK Biobank (UKB) | 92,693,895 | >0.001% | none | SNP + age + sex + array + PC1 + ... + PC10 | 72,355,667<sup>b</sup> | UKBB PCs | PLINK | 7,852,376 | 1.07 |
| TwinsUK | 27,445,314 | > 0% | ≥0.3 | SNP + age + sex + kinship (Unrelated individuals only) | 16,158,721 | GenABEL kinship matrix | R/GenABEL | 6,902,696 | 1.01 |
| Replication |  |  |  |  |  |  |  |  |  |
| UK Biobank (UKB) | 92,693,895 | >0.001% | none | SNP + age + sex + array + PC1 + ... + PC10 | 4<sup>bc</sup> | UKBB PCs | PLINK | NA | NA |

GWAS=genome-wide association study
<sup>a</sup>Meta-analysis LD score regression (LDSr) intercept was 1.007, indicating no population stratification. The LDSr intercept was used as a correction factor.
<sup>b</sup>only HRC-imputed SNPs were included in subsequent stages of analysis, including meta-analysis and replication, due to known problems at this time with the second set of imputed SNPs using 1000G + UK10K http://www.ukbiobank.ac.uk/2017/07/important-note-about-imputed-genetics-data/
<sup>c</sup>region-specific analyses were performed, and 4 variants were carried forward into the replication phase

**Supplemental Table S4:**
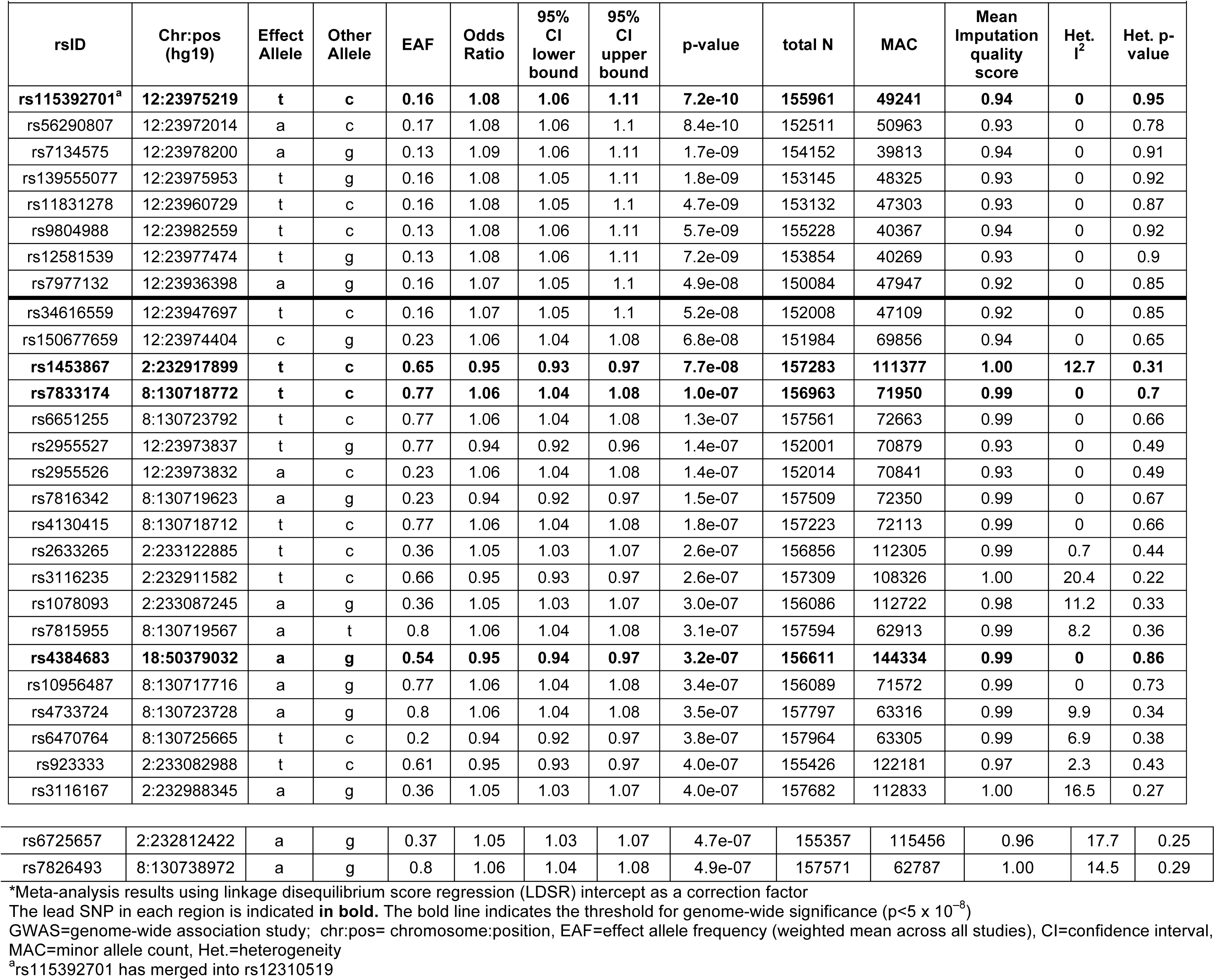
Association Results for Meta-Analysis of Chronic Back Pain GWAS (all variants with p<5 × 10^−7^)^*^

| rsID | Chr:pos<br>(hg19) | Effect<br>Allele | Other<br>Allele | EAF | Odds<br>Ratio | 95%<br>CI<br>lower<br>bound | 95%<br>CI<br>upper<br>bound | p-value | total N | MAC | Mean<br>Imputation<br>quality<br>score | Het.<br>$I^2$ | Het. p-value |
| --- | --- | --- | --- | --- | --- | --- | --- | --- | --- | --- | --- | --- | --- |
| **rs115392701<sup>a</sup>** | **12:23975219** | **t** | **c** | **0.16** | **1.08** | **1.06** | **1.11** | **7.2e-10** | **155961** | **49241** | **0.94** | **0** | **0.95** |
| rs56290807 | 12:23972014 | a | c | 0.17 | 1.08 | 1.06 | 1.1 | 8.4e-10 | 152511 | 50963 | 0.93 | 0 | 0.78 |
| rs7134575 | 12:23978200 | a | g | 0.13 | 1.09 | 1.06 | 1.11 | 1.7e-09 | 154152 | 39813 | 0.94 | 0 | 0.91 |
| rs139555077 | 12:23975953 | t | g | 0.16 | 1.08 | 1.05 | 1.11 | 1.8e-09 | 153145 | 48325 | 0.93 | 0 | 0.92 |
| rs11831278 | 12:23960729 | t | c | 0.16 | 1.08 | 1.05 | 1.1 | 4.7e-09 | 153132 | 47303 | 0.93 | 0 | 0.87 |
| rs9804988 | 12:23982559 | t | c | 0.13 | 1.08 | 1.06 | 1.11 | 5.7e-09 | 155228 | 40367 | 0.94 | 0 | 0.92 |
| rs12581539 | 12:23977474 | t | g | 0.13 | 1.08 | 1.06 | 1.11 | 7.2e-09 | 153854 | 40269 | 0.93 | 0 | 0.9 |
| rs7977132 | 12:23936398 | a | g | 0.16 | 1.07 | 1.05 | 1.1 | 4.9e-08 | 150084 | 47947 | 0.92 | 0 | 0.85 |
| rs34616559 | 12:23947697 | t | c | 0.16 | 1.07 | 1.05 | 1.1 | 5.2e-08 | 152008 | 47109 | 0.92 | 0 | 0.85 |
| rs150677659 | 12:23974404 | c | g | 0.23 | 1.06 | 1.04 | 1.08 | 6.8e-08 | 151984 | 69856 | 0.94 | 0 | 0.65 |
| **rs1453867** | **2:232917899** | **t** | **c** | **0.65** | **0.95** | **0.93** | **0.97** | **7.7e-08** | **157283** | **111377** | **1.00** | **12.7** | **0.31** |
| **rs7833174** | **8:130718772** | **t** | **c** | **0.77** | **1.06** | **1.04** | **1.08** | **1.0e-07** | **156963** | **71950** | **0.99** | **0** | **0.7** |
| rs6651255 | 8:130723792 | t | c | 0.77 | 1.06 | 1.04 | 1.08 | 1.3e-07 | 157561 | 72663 | 0.99 | 0 | 0.66 |
| rs2955527 | 12:23973837 | t | g | 0.77 | 0.94 | 0.92 | 0.96 | 1.4e-07 | 152001 | 70879 | 0.93 | 0 | 0.49 |
| rs2955526 | 12:23973832 | a | c | 0.23 | 1.06 | 1.04 | 1.08 | 1.4e-07 | 152014 | 70841 | 0.93 | 0 | 0.49 |
| rs7816342 | 8:130719623 | a | g | 0.23 | 0.94 | 0.92 | 0.97 | 1.5e-07 | 157509 | 72350 | 0.99 | 0 | 0.67 |
| rs4130415 | 8:130718712 | t | c | 0.77 | 1.06 | 1.04 | 1.08 | 1.8e-07 | 157223 | 72113 | 0.99 | 0 | 0.66 |
| rs2633265 | 2:233122885 | t | c | 0.36 | 1.05 | 1.03 | 1.07 | 2.6e-07 | 156856 | 112305 | 0.99 | 0.7 | 0.44 |
| rs3116235 | 2:232911582 | t | c | 0.66 | 0.95 | 0.93 | 0.97 | 2.6e-07 | 157309 | 108326 | 1.00 | 20.4 | 0.22 |
| rs1078093 | 2:233087245 | a | g | 0.36 | 1.05 | 1.03 | 1.07 | 3.0e-07 | 156086 | 112722 | 0.98 | 11.2 | 0.33 |
| rs7815955 | 8:130719567 | a | t | 0.8 | 1.06 | 1.04 | 1.08 | 3.1e-07 | 157594 | 62913 | 0.99 | 8.2 | 0.36 |
| **rs4384683** | **18:50379032** | **a** | **g** | **0.54** | **0.95** | **0.94** | **0.97** | **3.2e-07** | **156611** | **144334** | **0.99** | **0** | **0.86** |
| rs10956487 | 8:130717716 | a | g | 0.77 | 1.06 | 1.04 | 1.08 | 3.4e-07 | 156089 | 71572 | 0.99 | 0 | 0.73 |
| rs4733724 | 8:130723728 | a | g | 0.8 | 1.06 | 1.04 | 1.08 | 3.5e-07 | 157797 | 63316 | 0.99 | 9.9 | 0.34 |
| rs6470764 | 8:130725665 | t | c | 0.2 | 0.94 | 0.92 | 0.97 | 3.8e-07 | 157964 | 63305 | 0.99 | 6.9 | 0.38 |
| rs923333 | 2:233082988 | t | c | 0.61 | 0.95 | 0.93 | 0.97 | 4.0e-07 | 155426 | 122181 | 0.97 | 2.3 | 0.43 |
| rs3116167 | 2:232988345 | a | g | 0.36 | 1.05 | 1.03 | 1.07 | 4.0e-07 | 157682 | 112833 | 1.00 | 16.5 | 0.27 |
| rs6725657 | 2:232812422 | a | g | 0.37 | 1.05 | 1.03 | 1.07 | 4.7e-07 | 155357 | 115456 | 0.96 | 17.7 | 0.25 |
| rs7826493 | 8:130738972 | a | g | 0.8 | 1.06 | 1.04 | 1.08 | 4.9e-07 | 157571 | 62787 | 1.00 | 14.5 | 0.29 |

\*Meta-analysis results using linkage disequilibrium score regression (LDSR) intercept as a correction factor
The lead SNP in each region is indicated **in bold**. The bold line indicates the threshold for genome-wide significance ( $p < 5 \times 10^{-8}$ )
GWAS=genome-wide association study; chr:pos= chromosome:position, EAF=effect allele frequency (weighted mean across all studies), CI=confidence interval,
MAC=minor allele count, Het.=heterogeneity
<sup>a</sup>rs115392701 has merged into rs12310519

**Supplemental Table S5.** Variants associated with chronic back pain at the suggestive significance level (or proxy variants in LD [*r*^2^>0.6]), and associations with other phenotypes*

| rsID | Chr:pos (hg19) | Alleles | Proxy rsID | Proxy Pos (hg19) | Proxy Alleles (Effect/ Other) | $r^2$ | Trait | EAF | Beta | SE | P-value | N | N Studies | Unit | PMID |
| --- | --- | --- | --- | --- | --- | --- | --- | --- | --- | --- | --- | --- | --- | --- | --- |
| **SOX5** |  |  |  |  |  |  |  |  |  |  |  |  |  |  |  |
| rs115392701 / rs12310519 | chr12:23975219 | C/T | rs9804988 | chr12:23982559 | C/T | 0.727 | Height* | 0.90 | -0.024 | 0.0053 | 9.60E-06 | 236741 | 79 | Z-score | 25282103 |
| **CCDC26/GSDMC** |  |  |  |  |  |  |  |  |  |  |  |  |  |  |  |
| rs7833174 | chr8:130718772 | C/T | rs4733724 | chr8:130723728 | G/A | 0.845 | Height | 0.20 | -0.050 | 0.0037 | 1.00E-41 | NA | NA | unit increase | 25282103 |
| rs7833174 | chr8:130718772 | C/T | rs4733724 | chr8:130723728 | G/A | 0.845 | Height | 0.19 | -0.050 | 0.0037 | 1.40E-41 | 252637 | 79 | Z-score | 25282103 |
| rs7833174 | chr8:130718772 | C/T | rs7815955 | chr8:130719567 | T/A | 0.845 | Height | 0.20 | -0.050 | 0.0037 | 1.60E-41 | 252535 | 79 | Z-score | 25282103 |
| rs7833174 | chr8:130718772 | C/T | rs6470764 | chr8:130725665 | T/C | 0.845 | Height | 0.20 | -0.050 | 0.0037 | 4.30E-41 | 252663 | 79 | Z-score | 25282103 |
| rs7833174 | chr8:130718772 | C/T | rs4130415 | chr8:130718712 | C/T | 1 | Height | 0.19 | -0.050 | 0.0037 | 5.80E-41 | 252591 | 79 | Z-score | 25282103 |
| rs7833174 | chr8:130718772 | C/T | rs7816342 | chr8:130719623 | A/G | 1 | Height | 0.20 | -0.047 | 0.0036 | 2.00E-39 | 252715 | 79 | Z-score | 25282103 |
| rs7833174 | chr8:130718772 | C/T | rs2062078 | chr8:130734461 | T/G | 0.835 | Height | 0.20 | -0.047 | 0.0036 | 1.20E-38 | 253131 | 79 | Z-score | 25282103 |
| rs7833174 | chr8:130718772 | C/T | rs6470765 | chr8:130736697 | C/A | 0.835 | Height | 0.20 | -0.047 | 0.0036 | 1.40E-38 | 253054 | 79 | Z-score | 25282103 |
| rs7833174 | chr8:130718772 | C/T | rs6986256 | chr8:130728361 | A/T | 0.835 | Height | 0.20 | -0.047 | 0.0036 | 1.70E-38 | 253160 | 79 | Z-score | 25282103 |
| rs7833174 | chr8:130718772 | C/T | rs6470763 | chr8:130720646 | C/G | 0.737 | Height | 0.18 | -0.049 | 0.0038 | 2.90E-38 | 252464 | 79 | Z-score | 25282103 |
| rs7833174 | chr8:130718772 | C/T | rs3886937 | chr8:130737391 | C/T | 0.835 | Height | 0.20 | -0.047 | 0.0036 | 3.00E-38 | 253173 | 79 | Z-score | 25282103 |
| rs7833174 | chr8:130718772 | C/T | rs7833174 | chr8:130718772 | C/T | 1 | Height | 0.23 | -0.047 | 0.0036 | 4.50E-38 | 251729 | 79 | Z-score | 25282103 |
| rs7833174 | chr8:130718772 | C/T | rs4733732 | chr8:130738457 | T/C | 0.835 | Height | 0.20 | -0.047 | 0.0036 | 4.60E-38 | 253172 | 79 | Z-score | 25282103 |
| rs7833174 | chr8:130718772 | C/T | rs1074286 | chr8:130737601 | T/C | 0.835 | Height | 0.20 | -0.047 | 0.0036 | 5.10E-38 | 253173 | 79 | Z-score | 25282103 |
| rs7833174 | chr8:130718772 | C/T | rs3886938 | chr8:130737546 | G/T | 0.835 | Height | 0.20 | -0.047 | 0.0036 | 2.60E-37 | 252143 | 79 | Z-score | 25282103 |
| rs7833174 | chr8:130718772 | C/T | rs6985118 | chr8:130736929 | T/A | 0.835 | Height | 0.21 | -0.046 | 0.0037 | 1.10E-35 | 245733 | 79 | Z-score | 25282103 |
| rs7833174 | chr8:130718772 | C/T | rs4128895 | chr8:130755205 | C/T | 0.705 | Height | 0.18 | -0.044 | 0.0039 | 4.60E-29 | 253155 | 79 | Z-score | 25282103 |
| rs7833174 | chr8:130718772 | C/T | rs4733737 | chr8:130754719 | T/C | 0.71 | Height | 0.18 | -0.044 | 0.0039 | 5.00E-29 | 253204 | 79 | Z-score | 25282103 |
| rs7833174 | chr8:130718772 | C/T | rs7839339 | chr8:130743547 | A/T | 0.71 | Height | 0.18 | -0.044 | 0.0039 | 7.40E-29 | 253151 | 79 | Z-score | 25282103 |
| rs7833174 | chr8:130718772 | C/T | rs4236748 | chr8:130754887 | T/G | 0.71 | Height | 0.18 | -0.044 | 0.0039 | 8.00E-29 | 253158 | 79 | Z-score | 25282103 |
| rs7833174 | chr8:130718772 | C/T | rs1989396 | chr8:130756531 | A/G | 0.71 | Height | 0.18 | -0.044 | 0.0039 | 8.10E-29 | 253078 | 79 | Z-score | 25282103 |
| rs7833174 | chr8:130718772 | C/T | rs6994279 | chr8:130740650 | C/T | 0.71 | Height | 0.18 | -0.044 | 0.0040 | 1.30E-28 | 253160 | 79 | Z-score | 25282103 |
| rs7833174 | chr8:130718772 | C/T | rs7012249 | chr8:130740522 | A/G | 0.71 | Height | 0.18 | -0.044 | 0.0040 | 1.40E-28 | 253160 | 79 | Z-score | 25282103 |
| rs7833174 | chr8:130718772 | C/T | rs6998296 | chr8:130740779 | C/T | 0.71 | Height | 0.18 | -0.044 | 0.0040 | 1.40E-28 | 253160 | 79 | Z-score | 25282103 |
| rs7833174 | chr8:130718772 | C/T | rs6470764 | chr8:130725665 | T/C | 0.845 | Height | NA | NA | NA | 1.70E-28 | 11513 | NA | NA | 23704328 |
| rs7833174 | chr8:130718772 | C/T | rs6470764 | chr8:130725665 | T/C | 0.845 | Height | NA | NA | NA | 1.70E-28 | 183727 | NA | NA | 20881960 |
| rs7833174 | chr8:130718772 | C/T | rs6470764 | chr8:130725665 | T/C | 0.845 | Height | NA | NA | NA | 1.70E-28 | 94612 | NA | NA | 21946350 |
| rs7833174 | chr8:130718772 | C/T | rs7011557 | chr8:130740135 | T/G | 0.71 | Height | 0.18 | -0.044 | 0.0040 | 1.90E-28 | 253160 | 79 | Z-score | 25282103 |
| **rs7833174** | **chr8:130718772** | **C/T** | **rs6470764** | **chr8:130725665** | **T/C** | **0.845** | **Height** | **0.20** | **0.050** | **0.0045** | **2.00E-28** | **NA** | **NA** | **unit decrease** | **20881960** |
| rs7833174 | chr8:130718772 | C/T | rs6470778 | chr8:130759579 | T/C | 0.71 | Height | 0.18 | -0.043 | 0.0039 | 2.60E-28 | 253090 | 79 | Z-score | 25282103 |
| rs7833174 | chr8:130718772 | C/T | rs6470768 | chr8:130739528 | A/G | 0.71 | Height | 0.18 | -0.044 | 0.0040 | 4.70E-28 | 252174 | 79 | Z-score | 25282103 |
| rs7833174 | chr8:130718772 | C/T | rs6470766 | chr8:130736854 | G/A | 0.706 | Height | 0.18 | -0.044 | 0.0040 | 1.30E-27 | 246945 | 79 | Z-score | 25282103 |
| **rs7833174** | **chr8:130718772** | **C/T** | **rs4733724** | **chr8:130723728** | **G/A** | **0.845** | **Height** | **0.19** | **-0.047** | **0.0053** | **6.57E-19** | **133673** | **46** | **Z-score** | **23754948** |
| **rs7833174** | **chr8:130718772** | **C/T** | **rs6470764** | **chr8:130725665** | **T/C** | **0.845** | **Height** | **0.19** | **-0.047** | **0.0053** | **6.57E-19** | **133673** | **46** | **Z-score** | **23754948** |
| **rs7833174** | **chr8:130718772** | **C/T** | **rs4130415** | **chr8:130718712** | **C/T** | **1** | **Height** | **0.19** | **-0.047** | **0.0053** | **1.60E-18** | **133671** | **46** | **Z-score** | **23754948** |
| **rs7833174** | **chr8:130718772** | **C/T** | **rs7815955** | **chr8:130719567** | **T/A** | **0.845** | **Height** | **0.19** | **-0.047** | **0.0053** | **1.60E-18** | **133673** | **46** | **Z-score** | **23754948** |
| rs7833174 | chr8:130718772 | C/T | rs6470765 | chr8:130736697 | C/A | 0.835 | Height | 0.20 | -0.044 | 0.0051 | 1.59E-17 | 133549 | 46 | Z-score | 23754948 |
| **rs7833174** | **chr8:130718772** | **C/T** | **rs7833174** | **chr8:130718772** | **C/T** | **1** | **Height** | **0.20** | **-0.043** | **0.0051** | **2.00E-17** | **133668** | **46** | **Z-score** | **23754948** |
| **rs7833174** | **chr8:130718772** | **C/T** | **rs7816342** | **chr8:130719623** | **A/G** | **1** | **Height** | **0.20** | **-0.043** | **0.0051** | **2.00E-17** | **133669** | **46** | **Z-score** | **23754948** |
| rs7833174 | chr8:130718772 | C/T | rs6470763 | chr8:130720646 | C/G | 0.737 | Height | 0.18 | -0.046 | 0.0054 | 2.25E-17 | 133683 | 46 | Z-score | 23754948 |
| rs7833174 | chr8:130718772 | C/T | rs6986256 | chr8:130728361 | A/T | 0.835 | Height | 0.20 | -0.044 | 0.0052 | 2.85E-17 | 133668 | 46 | Z-score | 23754948 |
| rs7833174 | chr8:130718772 | C/T | rs3886937 | chr8:130737391 | C/T | 0.835 | Height | 0.20 | -0.044 | 0.0052 | 2.85E-17 | 133669 | 46 | Z-score | 23754948 |
| rs7833174 | chr8:130718772 | C/T | rs1074286 | chr8:130737601 | T/C | 0.835 | Height | 0.20 | -0.044 | 0.0052 | 2.85E-17 | 133669 | 46 | Z-score | 23754948 |
| rs7833174 | chr8:130718772 | C/T | rs4733732 | chr8:130738457 | T/C | 0.835 | Height | 0.20 | -0.044 | 0.0052 | 2.85E-17 | 133667 | 46 | Z-score | 23754948 |
| rs7833174 | chr8:130718772 | C/T | rs2062078 | chr8:130734461 | T/G | 0.835 | Height | 0.20 | -0.043 | 0.0051 | 3.89E-17 | 133605 | 46 | Z-score | 23754948 |
| rs7833174 | chr8:130718772 | C/T | rs3886938 | chr8:130737546 | G/T | 0.835 | Height | 0.20 | -0.043 | 0.0051 | 3.89E-17 | 133628 | 46 | Z-score | 23754948 |
| **rs7833174** | **chr8:130718772** | **C/T** | **rs7815955** | **chr8:130719567** | **T/A** | **0.845** | **Height** | **NA** | **NA** | **NA** | **7.95E-17** | **183727** | **NA** | **NA** | **20881960** |
| **rs7833174** | **chr8:130718772** | **C/T** | **rs4733724** | **chr8:130723728** | **G/A** | **0.845** | **Height** | **NA** | **NA** | **NA** | **9.24E-17** | **183727** | **NA** | **NA** | **20881960** |
| rs7833174 | chr8:130718772 | C/T | rs6985118 | chr8:130736929 | T/A | 0.835 | Height | 0.21 | -0.043 | 0.0052 | 2.44E-16 | 131884 | 46 | Z-score | 23754948 |
| **rs7833174** | **chr8:130718772** | **C/T** | **rs4130415** | **chr8:130718712** | **C/T** | **1** | **Height** | **NA** | **NA** | **NA** | **2.54E-16** | **183727** | **NA** | **NA** | **20881960** |
| **rs7833174** | **chr8:130718772** | **C/T** | **rs7816342** | **chr8:130719623** | **A/G** | **1** | **Height** | **NA** | **NA** | **NA** | **1.41E-15** | **183727** | **NA** | **NA** | **20881960** |
| **rs7833174** | **chr8:130718772** | **C/T** | **rs7833174** | **chr8:130718772** | **C/T** | **1** | **Height** | **NA** | **NA** | **NA** | **1.64E-15** | **183727** | **NA** | **NA** | **20881960** |
| rs7833174 | chr8:130718772 | C/T | rs6470763 | chr8:130720646 | C/G | 0.737 | Height | NA | NA | NA | 1.64E-15 | 183727 | NA | NA | 20881960 |
| rs7833174 | chr8:130718772 | C/T | rs6470765 | chr8:130736697 | C/A | 0.835 | Height | NA | NA | NA | 1.90E-15 | 183727 | NA | NA | 20881960 |
| rs7833174 | chr8:130718772 | C/T | rs6986256 | chr8:130728361 | A/T | 0.835 | Height | NA | NA | NA | 2.20E-15 | 183727 | NA | NA | 20881960 |
| rs7833174 | chr8:130718772 | C/T | rs3886937 | chr8:130737391 | C/T | 0.835 | Height | NA | NA | NA | 2.94E-15 | 183727 | NA | NA | 20881960 |
| rs7833174 | chr8:130718772 | C/T | rs4733732 | chr8:130738457 | T/C | 0.835 | Height | NA | NA | NA | 2.94E-15 | 183727 | NA | NA | 20881960 |
| rs7833174 | chr8:130718772 | C/T | rs1074286 | chr8:130737601 | T/C | 0.835 | Height | NA | NA | NA | 4.55E-15 | 183727 | NA | NA | 20881960 |
| rs7833174 | chr8:130718772 | C/T | rs2062078 | chr8:130734461 | T/G | 0.835 | Height | NA | NA | NA | 5.26E-15 | 183727 | NA | NA | 20881960 |
| rs7833174 | chr8:130718772 | C/T | rs3886938 | chr8:130737546 | G/T | 0.835 | Height | NA | NA | NA | 5.26E-15 | 183727 | NA | NA | 20881960 |
| rs7833174 | chr8:130718772 | C/T | rs4559213 | chr8:130753742 | C/T | 0.71 | Height | 0.18 | -0.055 | 0.0070 | 8.60E-15 | 73853 | 79 | Z-score | 25282103 |
| rs7833174 | chr8:130718772 | C/T | rs4633026 | chr8:130752408 | C/T | 0.71 | Height | 0.18 | -0.055 | 0.0071 | 1.00E-14 | 73853 | 79 | Z-score | 25282103 |
| rs7833174 | chr8:130718772 | C/T | rs6985118 | chr8:130736929 | T/A | 0.835 | Height | NA | NA | NA | 1.66E-14 | 183727 | NA | NA | 20881960 |
| rs7833174 | chr8:130718772 | C/T | rs7839339 | chr8:130743547 | A/T | 0.71 | Height | 0.18 | -0.041 | 0.0056 | 1.14E-13 | 133666 | 46 | Z-score | 23754948 |
| rs7833174 | chr8:130718772 | C/T | rs4733737 | chr8:130754719 | T/C | 0.71 | Height | 0.18 | -0.041 | 0.0056 | 1.14E-13 | 133678 | 46 | Z-score | 23754948 |
| rs7833174 | chr8:130718772 | C/T | rs4236748 | chr8:130754887 | T/G | 0.71 | Height | 0.18 | -0.041 | 0.0056 | 1.14E-13 | 133678 | 46 | Z-score | 23754948 |
| rs7833174 | chr8:130718772 | C/T | rs4128895 | chr8:130755205 | C/T | 0.705 | Height | 0.18 | -0.041 | 0.0056 | 1.14E-13 | 133678 | 46 | Z-score | 23754948 |
| rs7833174 | chr8:130718772 | C/T | rs6470768 | chr8:130739528 | A/G | 0.71 | Height | 0.18 | -0.041 | 0.0056 | 2.35E-13 | 133677 | 46 | Z-score | 23754948 |
| rs7833174 | chr8:130718772 | C/T | rs7011557 | chr8:130740135 | T/G | 0.71 | Height | 0.18 | -0.041 | 0.0056 | 2.35E-13 | 133677 | 46 | Z-score | 23754948 |
| rs7833174 | chr8:130718772 | C/T | rs7012249 | chr8:130740522 | A/G | 0.71 | Height | 0.18 | -0.041 | 0.0056 | 2.35E-13 | 133676 | 46 | Z-score | 23754948 |
| rs7833174 | chr8:130718772 | C/T | rs6994279 | chr8:130740650 | C/T | 0.71 | Height | 0.18 | -0.041 | 0.0056 | 2.35E-13 | 133676 | 46 | Z-score | 23754948 |
| rs7833174 | chr8:130718772 | C/T | rs6998296 | chr8:130740779 | C/T | 0.71 | Height | 0.18 | -0.041 | 0.0056 | 2.35E-13 | 133677 | 46 | Z-score | 23754948 |
| rs7833174 | chr8:130718772 | C/T | rs1989396 | chr8:130756531 | A/G | 0.71 | Height | 0.18 | -0.041 | 0.0056 | 2.35E-13 | 133607 | 46 | Z-score | 23754948 |
| rs7833174 | chr8:130718772 | C/T | rs6470778 | chr8:130759579 | T/C | 0.71 | Height | 0.18 | -0.040 | 0.0055 | 2.71E-13 | 133621 | 46 | Z-score | 23754948 |
| rs7833174 | chr8:130718772 | C/T | rs6470766 | chr8:130736854 | G/A | 0.706 | Height | 0.18 | -0.041 | 0.0056 | 2.96E-13 | 133693 | 46 | Z-score | 23754948 |
| rs7833174 | chr8:130718772 | C/T | rs4733737 | chr8:130754719 | T/C | 0.71 | Height | NA | NA | NA | 6.92E-12 | 183727 | NA | NA | 20881960 |
| rs7833174 | chr8:130718772 | C/T | rs7839339 | chr8:130743547 | A/T | 0.71 | Height | NA | NA | NA | 7.78E-12 | 183727 | NA | NA | 20881960 |
| rs7833174 | chr8:130718772 | C/T | rs6470766 | chr8:130736854 | G/A | 0.706 | Height | NA | NA | NA | 7.78E-12 | 183727 | NA | NA | 20881960 |
| rs7833174 | chr8:130718772 | C/T | rs6470768 | chr8:130739528 | A/G | 0.71 | Height | NA | NA | NA | 1.10E-11 | 183727 | NA | NA | 20881960 |
| rs7833174 | chr8:130718772 | C/T | rs7011557 | chr8:130740135 | T/G | 0.71 | Height | NA | NA | NA | 1.10E-11 | 183727 | NA | NA | 20881960 |
| rs7833174 | chr8:130718772 | C/T | rs4236748 | chr8:130754887 | T/G | 0.71 | Height | NA | NA | NA | 1.10E-11 | 183727 | NA | NA | 20881960 |
| rs7833174 | chr8:130718772 | C/T | rs4128895 | chr8:130755205 | C/T | 0.705 | Height | NA | NA | NA | 1.24E-11 | 183727 | NA | NA | 20881960 |
| rs7833174 | chr8:130718772 | C/T | rs7012249 | chr8:130740522 | A/G | 0.71 | Height | NA | NA | NA | 1.39E-11 | 183727 | NA | NA | 20881960 |
| rs7833174 | chr8:130718772 | C/T | rs1989396 | chr8:130756531 | A/G | 0.71 | Height | NA | NA | NA | 1.39E-11 | 183727 | NA | NA | 20881960 |
| rs7833174 | chr8:130718772 | C/T | rs6994279 | chr8:130740650 | C/T | 0.71 | Height | NA | NA | NA | 1.56E-11 | 183727 | NA | NA | 20881960 |
| rs7833174 | chr8:130718772 | C/T | rs6998296 | chr8:130740779 | C/T | 0.71 | Height | NA | NA | NA | 1.56E-11 | 183727 | NA | NA | 20881960 |
| rs7833174 | chr8:130718772 | C/T | rs6470778 | chr8:130759579 | T/C | 0.71 | Height | NA | NA | NA | 1.75E-11 | 183727 | NA | NA | 20881960 |
| **rs7833174** | **chr8:130718772** | **C/T** | **rs4733724** | **chr8:130723728** | **G/A** | **0.845** | **Height in females** | **0.19** | **-0.048** | **0.0073** | **5.68E-11** | **73124** | **46** | **Z-score** | **23754948** |
| **rs7833174** | **chr8:130718772** | **C/T** | **rs6470764** | **chr8:130725665** | **T/C** | **0.845** | **Height in females** | **0.19** | **-0.048** | **0.0073** | **7.47E-11** | **73124** | **46** | **Z-score** | **23754948** |
| **rs7833174** | **chr8:130718772** | **C/T** | **rs4130415** | **chr8:130718712** | **C/T** | **1** | **Height in females** | **0.19** | **-0.047** | **0.0073** | **8.96E-11** | **73124** | **46** | **Z-score** | **23754948** |
| **rs7833174** | **chr8:130718772** | **C/T** | **rs7815955** | **chr8:130719567** | **T/A** | **0.845** | **Height in females** | **0.19** | **-0.047** | **0.0073** | **8.96E-11** | **73124** | **46** | **Z-score** | **23754948** |
| rs7833174 | chr8:130718772 | C/T | rs6470765 | chr8:130736697 | C/A | 0.835 | Height in females | 0.20 | -0.045 | 0.0070 | 9.06E-11 | 73054 | 46 | Z-score | 23754948 |
| rs7833174 | chr8:130718772 | C/T | rs3886938 | chr8:130737546 | G/T | 0.835 | Height in females | 0.20 | -0.044 | 0.0070 | 1.93E-10 | 73100 | 46 | Z-score | 23754948 |
| rs7833174 | chr8:130718772 | C/T | rs2062078 | chr8:130734461 | T/G | 0.835 | Height in females | 0.20 | -0.044 | 0.0070 | 2.11E-10 | 73082 | 46 | Z-score | 23754948 |
| rs7833174 | chr8:130718772 | C/T | rs4733732 | chr8:130738457 | T/C | 0.835 | Height in females | 0.20 | -0.045 | 0.0071 | 2.82E-10 | 73122 | 46 | Z-score | 23754948 |
| rs7833174 | chr8:130718772 | C/T | rs6986256 | chr8:130728361 | A/T | 0.835 | Height in females | 0.20 | -0.045 | 0.0071 | 3.09E-10 | 73123 | 46 | Z-score | 23754948 |
| rs7833174 | chr8:130718772 | C/T | rs3886937 | chr8:130737391 | C/T | 0.835 | Height in females | 0.20 | -0.045 | 0.0071 | 3.09E-10 | 73124 | 46 | Z-score | 23754948 |
| rs7833174 | chr8:130718772 | C/T | rs1074286 | chr8:130737601 | T/C | 0.835 | Height in females | 0.20 | -0.045 | 0.0071 | 3.09E-10 | 73124 | 46 | Z-score | 23754948 |
| rs7833174 | chr8:130718772 | C/T | rs6470763 | chr8:130720646 | C/G | 0.737 | Height in females | 0.18 | -0.046 | 0.0074 | 4.09E-10 | 73129 | 46 | Z-score | 23754948 |
| **rs7833174** | **chr8:130718772** | **C/T** | **rs7833174** | **chr8:130718772** | **C/T** | **1** | **Height in females** | **0.20** | **-0.043** | **0.0070** | **6.38E-10** | **73124** | **46** | **Z-score** | **23754948** |
| **rs7833174** | **chr8:130718772** | **C/T** | **rs7816342** | **chr8:130719623** | **A/G** | **1** | **Height in females** | **0.20** | **-0.043** | **0.0070** | **6.38E-10** | **73124** | **46** | **Z-score** | **23754948** |
| rs7833174 | chr8:130718772 | C/T | rs4633026 | chr8:130752408 | C/T | 0.71 | Height | NA | -0.050 | 0.0081 | 9.63E-10 | 57807 | 46 | Z-score | 23754948 |
| rs7833174 | chr8:130718772 | C/T | rs4559213 | chr8:130753742 | C/T | 0.71 | Height | NA | -0.050 | 0.0081 | 9.63E-10 | 57807 | 46 | Z-score | 23754948 |
| rs7833174 | chr8:130718772 | C/T | rs6985118 | chr8:130736929 | T/A | 0.835 | Height in females | 0.21 | -0.043 | 0.0071 | 9.91E-10 | 72209 | 46 | Z-score | 23754948 |
| rs7833174 | chr8:130718772 | C/T | rs6470778 | chr8:130759579 | T/C | 0.71 | Height in females | 0.18 | -0.045 | 0.0075 | 1.84E-09 | 73097 | 46 | Z-score | 23754948 |
| **rs7833174** | **chr8:130718772** | **C/T** | **rs4130415** | **chr8:130718712** | **C/T** | **1** | **Height in males** | **0.19** | **-0.046** | **0.0077** | **1.95E-09** | **60547** | **46** | **Z-score** | **23754948** |
| **rs7833174** | **chr8:130718772** | **C/T** | **rs7815955** | **chr8:130719567** | **T/A** | **0.845** | **Height in males** | **0.19** | **-0.046** | **0.0077** | **2.11E-09** | **60549** | **46** | **Z-score** | **23754948** |
| rs7833174 | chr8:130718772 | C/T | rs4733737 | chr8:130754719 | T/C | 0.71 | Height in females | 0.18 | -0.046 | 0.0076 | 2.12E-09 | 73126 | 46 | Z-score | 23754948 |
| rs7833174 | chr8:130718772 | C/T | rs4236748 | chr8:130754887 | T/G | 0.71 | Height in females | 0.18 | -0.046 | 0.0076 | 2.30E-09 | 73126 | 46 | Z-score | 23754948 |
| rs7833174 | chr8:130718772 | C/T | rs7839339 | chr8:130743547 | A/T | 0.71 | Height in females | 0.18 | -0.046 | 0.0076 | 2.70E-09 | 73118 | 46 | Z-score | 23754948 |
| rs7833174 | chr8:130718772 | C/T | rs4128895 | chr8:130755205 | C/T | 0.705 | Height in females | 0.18 | -0.046 | 0.0076 | 2.70E-09 | 73126 | 46 | Z-score | 23754948 |
| **rs7833174** | **chr8:130718772** | **C/T** | **rs6470764** | **chr8:130725665** | **T/C** | **0.845** | **Height in males** | **0.19** | **-0.046** | **0.0077** | **2.90E-09** | **60549** | **46** | **Z-score** | **23754948** |
| **rs7833174** | **chr8:130718772** | **C/T** | **rs4733724** | **chr8:130723728** | **G/A** | **0.845** | **Height in males** | **0.19** | **-0.046** | **0.0077** | **3.14E-09** | **60549** | **46** | **Z-score** | **23754948** |
| rs7833174 | chr8:130718772 | C/T | rs7011557 | chr8:130740135 | T/G | 0.71 | Height in females | 0.18 | -0.045 | 0.0076 | 4.01E-09 | 73124 | 46 | Z-score | 23754948 |
| rs7833174 | chr8:130718772 | C/T | rs1989396 | chr8:130756531 | A/G | 0.71 | Height in females | 0.18 | -0.045 | 0.0076 | 4.01E-09 | 73088 | 46 | Z-score | 23754948 |
| rs7833174 | chr8:130718772 | C/T | rs6470766 | chr8:130736854 | G/A | 0.706 | Height in females | 0.18 | -0.045 | 0.0076 | 4.01E-09 | 73136 | 46 | Z-score | 23754948 |
| rs7833174 | chr8:130718772 | C/T | rs6470768 | chr8:130739528 | A/G | 0.71 | Height in females | 0.18 | -0.045 | 0.0076 | 4.34E-09 | 73125 | 46 | Z-score | 23754948 |
| **rs7833174** | **chr8:130718772** | **C/T** | **rs7816342** | **chr8:130719623** | **A/G** | **1** | **Height in males** | **0.20** | **-0.044** | **0.0075** | **4.61E-09** | **60545** | **46** | **Z-score** | **23754948** |
| rs7833174 | chr8:130718772 | C/T | rs7012249 | chr8:130740522 | A/G | 0.71 | Height in females | 0.18 | -0.045 | 0.0076 | 4.70E-09 | 73124 | 46 | Z-score | 23754948 |
| rs7833174 | chr8:130718772 | C/T | rs6994279 | chr8:130740650 | C/T | 0.71 | Height in females | 0.18 | -0.045 | 0.0076 | 4.70E-09 | 73124 | 46 | Z-score | 23754948 |
| rs7833174 | chr8:130718772 | C/T | rs6998296 | chr8:130740779 | C/T | 0.71 | Height in females | 0.18 | -0.045 | 0.0076 | 5.08E-09 | 73124 | 46 | Z-score | 23754948 |
| **rs7833174** | **chr8:130718772** | **C/T** | **rs7833174** | **chr8:130718772** | **C/T** | **1** | **Height in males** | **0.20** | **-0.044** | **0.0075** | **5.42E-09** | **60544** | **46** | **Z-score** | **23754948** |
| rs7833174 | chr8:130718772 | C/T | rs6470763 | chr8:130720646 | C/G | 0.737 | Height in males | 0.18 | -0.045 | 0.0078 | 8.28E-09 | 60554 | 46 | Z-score | 23754948 |
| rs7833174 | chr8:130718772 | C/T | rs6986256 | chr8:130728361 | A/T | 0.835 | Height in males | 0.20 | -0.042 | 0.0075 | 1.91E-08 | 60545 | 46 | Z-score | 23754948 |
| rs7833174 | chr8:130718772 | C/T | rs2062078 | chr8:130734461 | T/G | 0.835 | Height in males | 0.20 | -0.042 | 0.0075 | 2.06E-08 | 60523 | 46 | Z-score | 23754948 |
| rs7833174 | chr8:130718772 | C/T | rs4733732 | chr8:130738457 | T/C | 0.835 | Height in males | 0.20 | -0.042 | 0.0075 | 2.06E-08 | 60545 | 46 | Z-score | 23754948 |
| rs7833174 | chr8:130718772 | C/T | rs1074286 | chr8:130737601 | T/C | 0.835 | Height in males | 0.20 | -0.042 | 0.0075 | 2.23E-08 | 60545 | 46 | Z-score | 23754948 |
| rs7833174 | chr8:130718772 | C/T | rs3886937 | chr8:130737391 | C/T | 0.835 | Height in males | 0.20 | -0.042 | 0.0075 | 2.40E-08 | 60545 | 46 | Z-score | 23754948 |
| rs7833174 | chr8:130718772 | C/T | rs3886938 | chr8:130737546 | G/T | 0.835 | Height in males | 0.20 | -0.042 | 0.0075 | 2.40E-08 | 60528 | 46 | Z-score | 23754948 |
| rs7833174 | chr8:130718772 | C/T | rs6470765 | chr8:130736697 | C/A | 0.835 | Height in males | 0.20 | -0.042 | 0.0075 | 2.59E-08 | 60495 | 46 | Z-score | 23754948 |
| rs7833174 | chr8:130718772 | C/T | rs6985118 | chr8:130736929 | T/A | 0.835 | Height in males | 0.21 | -0.042 | 0.0076 | 4.14E-08 | 59676 | 46 | Z-score | 23754948 |
| rs7833174 | chr8:130718772 | C/T | rs4633026 | chr8:130752408 | C/T | 0.71 | Height in females | NA | -0.061 | 0.0120 | 2.00E-07 | 28350 | 46 | Z-score | 23754948 |
| rs7833174 | chr8:130718772 | C/T | rs4559213 | chr8:130753742 | C/T | 0.71 | Height in females | NA | -0.061 | 0.0120 | 2.00E-07 | 28350 | 46 | Z-score | 23754948 |
| **rs7833174** | **chr8:130718772** | **C/T** | **rs6470764** | **chr8:130725665** | **T/C** | **0.845** | **Infant length** | **0.20** | **0.054** | **0.0110** | **1.00E-06** | **NA** | **NA** | **unit decrease** | **25281659** |
| rs7833174 | chr8:130718772 | C/T | rs4733737 | chr8:130754719 | T/C | 0.71 | Height in males | 0.18 | -0.036 | 0.0082 | 8.20E-06 | 60552 | 46 | Z-score | 23754948 |
| rs7833174 | chr8:130718772 | C/T | rs6470768 | chr8:130739528 | A/G | 0.71 | Height in males | 0.18 | -0.036 | 0.0082 | 9.19E-06 | 60552 | 46 | Z-score | 23754948 |

**Bolded lines** indicate variants associated with chronic back pain in the discovery stage meta-analysis that were also associated with other phenotypes
Chr:pos=chromosome:position, EAF=effect allele frequency, SE=standard error, Het.= heterogeneity
^Variants associated with back pain at suggestive or genome-wide significance level ( $p < 5.0 \times 10^{-7}$ ) in the discovery stage, which achieved genome-wide significance ( $p < 5.0 \times 10^{-8}$ ) in the joint meta-analysis (discovery-replication), and which were also associated with any other phenotypes as identified by PhenoScanner, download date 11/22/2017. Associations with other phenotypes listed in this table are limited to those with $p < 5 \times 10^{-5}$ . All results are for Europeans.

**Supplemental Table S6.**
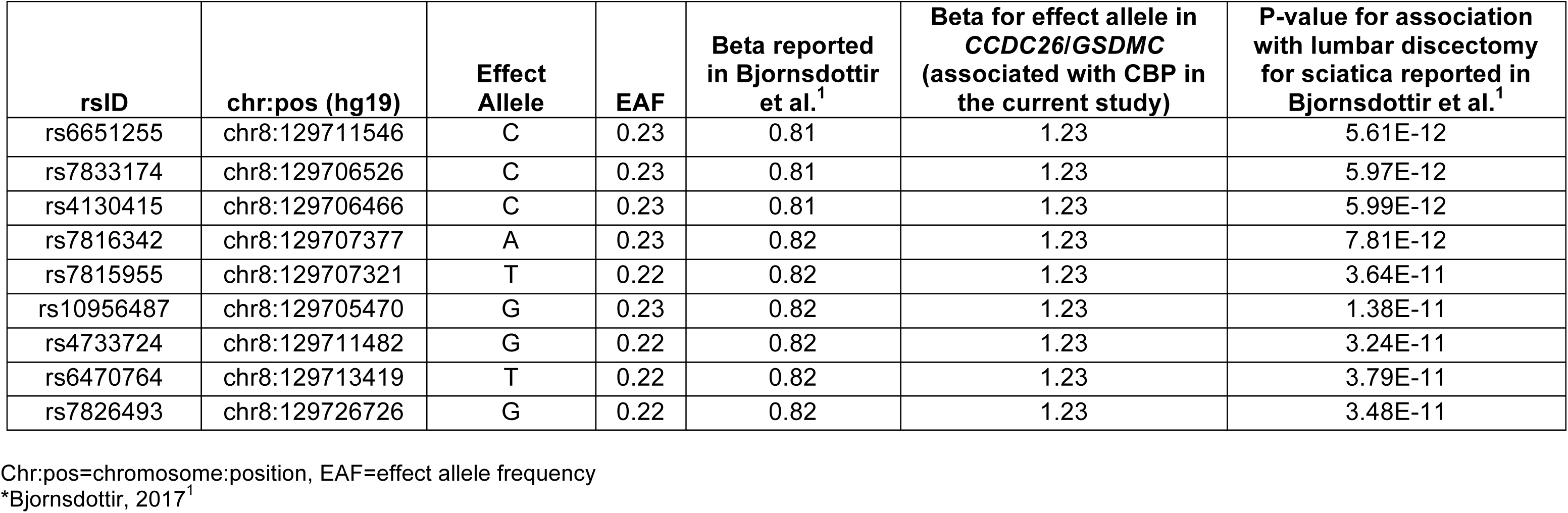
Variants in *CCDC26/GSDMC* associated with chronic back pain at the suggestive significance level (p<5 × 10^−7^) in the discovery stage meta-analysis, and associations with lumbar discectomy for sciatica in a prior GWAS*

**Supplemental Table S7:** Region-specific secondary analyses accounting for height, conducted in the UKB interim data release sample*

|  |  |  |  |  |  |  |  |  |  |  |  |  |
| --- | --- | --- | --- | --- | --- | --- | --- | --- | --- | --- | --- | --- |
| Associations with CBP (for the CBP-associated lead SNPs in SOX5 and CCDC26/GSDMC from the discovery stage meta-analysis), with and without adjustment for height as a covariate |  |  |  |  |  |  |  |  |  |  |  |  |
|  |  |  |  |  |  | Model 1 |  |  | Model 2 |  |  |  |
| rsID | location/gene | chr:pos (hg19) | Effect Allele | Other Allele | EAF | N | Beta | p-value | N | Beta | p-value |  |
| rs12310519 | intronic/SOX5 | 12:23975219 | T | C | 0.16 | 118108 | 0.076 | 1.32E-07 | 117917 | 1.11 | 7.2e-10 |  |
| rs7833174 | intergenic<br>CCDC26/GSDMC | 8:130718772 | T | C | 0.77 | 119007 | 0.060 | 2.71E-06 | 118815 | 1.10 | 8.4e-10 |  |
| Associations with CBP (for the CBP-associated lead SNPs in SOX5 and CCDC26/GSDMC from region-specific analyses in UKB interim data release sample), with and without conditioning on the top height-associated variant in the region |  |  |  |  |  |  |  |  |  |  |  |  |
|  |  |  |  |  |  | Model 1 |  |  | Model 3 |  |  |  |
| rsID | location/gene | chr:pos (hg19) | Effect Allele | Other Allele | EAF | N | Beta | p-value | N | Beta | p-value | conditional rsID<sup>a</sup> |
| rs7134575 | intronic/SOX5 | 12:23978200 | A | G | 0.13 | 118947 | 0.087 | 2.00E-08 | 117309 | 0.087 | 1.97E-08 | rs1498873 |
| rs7833174 | intergenic/<br>CCDC26/GSDMC | 8:130718772 | C | T | 0.23 | 119007 | -0.060 | 2.71E-06 | 118674 | -0.055 | 0.11 | rs3886937 |
| Associations with height (for the CBP-associated lead SNPs in SOX5 and CCDC26/GSDMC from region-specific analyses in UKB interim data release sample), with and without conditioning on the top CBP-associated variant in the region |  |  |  |  |  |  |  |  |  |  |  |  |
|  |  |  |  |  |  | Model 4 |  |  | Model 5 |  |  |  |
| rsID | location/gene | chr:pos (hg19) | Effect Allele | Other Allele | EAF | N | Beta | p-value | N | Beta | p-value | conditional rsID<sup>b</sup> |
| rs1498873 | intronic/SOX5 | 12:24200700 | T | G | 0.22 | 118181 | 0.200 | 1.38E-10 | 117125 | 0.19 | 4.36E-10 | rs7134575 |
| rs3886937 | intergenic/<br>CCDC26/GSDMC | 8:130737391 | C | T | 0.20 | 119479 | -0.373 | 6.79E-31 | 118482 | -0.251 | 0.0030 | rs7833174 |

UKB= UK biobank, CBP=chronic back pain, chr:pos=chromosome:position, EAF=effect allele frequency
\*all results reflect analyses conducted in the UKB interim data release sample only
<sup>a</sup>conditional on the top height-associated SNP in the region
<sup>a</sup>conditional on the top CBP-associated SNP in the region
Model 1: CBP ~ SNP + age + sex + array + PC1 + ... + PC10
Model 2: CBP ~ SNP + age + sex + height + array + PC1 + ... + PC10
Model 3: CBP ~ SNP + age + sex + array + PC1 + ... + PC10 (conditional on the top height-associated variant in the region)
Model 4: height ~ SNP + age + sex + array + PC1 + ... + PC10
Model 5: height ~ SNP + age + sex + array + PC1 + ... + PC10 (conditional on the top CBP-associated variant in the region)

**Supplemental Table S8.**
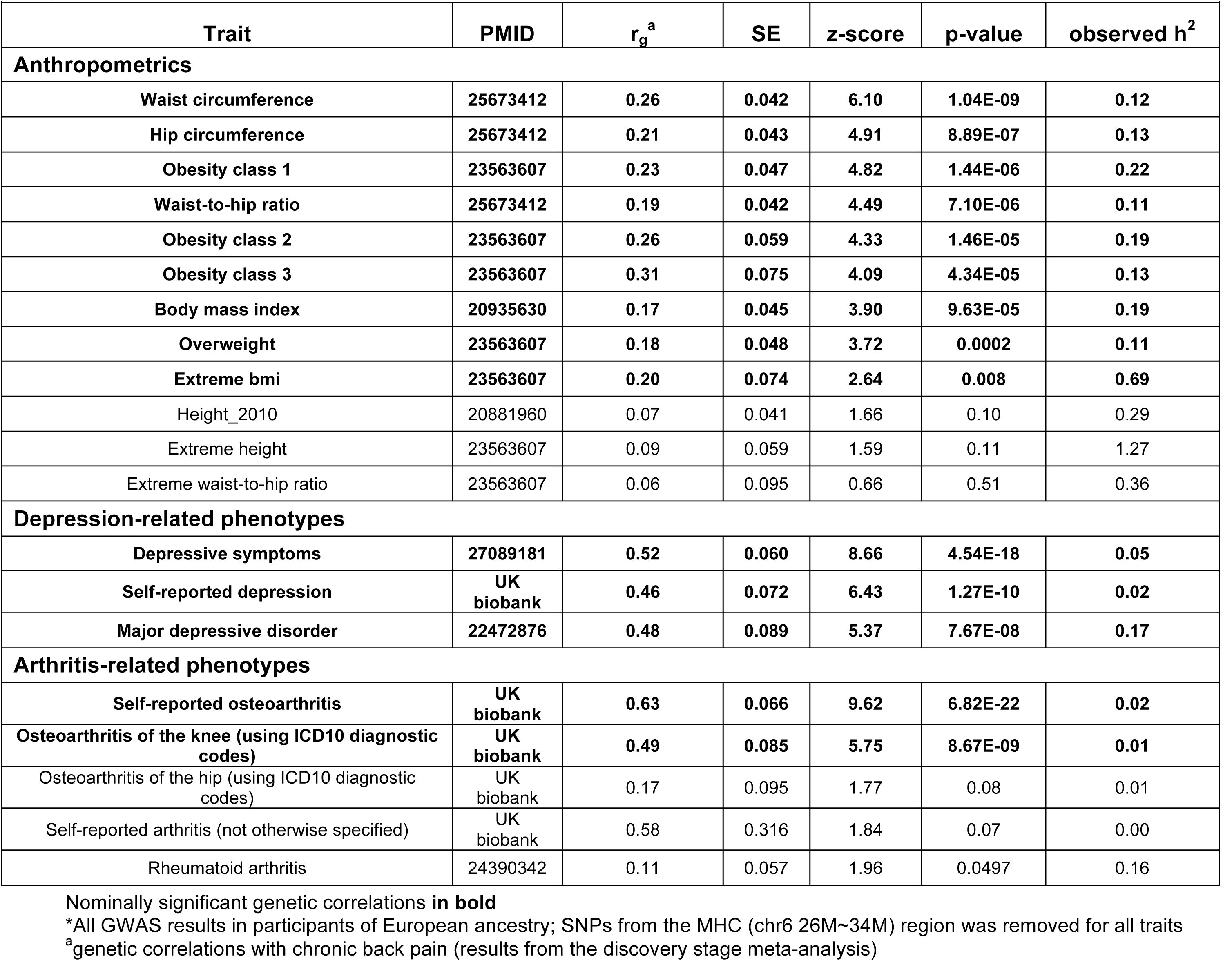
1. Genetic correlations between CBP and selected phenotypes of conceptual relevance to CBP, using cross-trait LD score regression*

| Trait | PMID | $r_g^a$ | SE | z-score | p-value | observed $h^2$ |
| --- | --- | --- | --- | --- | --- | --- |
| **Anthropometrics** |  |  |  |  |  |  |
| Waist circumference | 25673412 | 0.26 | 0.042 | 6.10 | 1.04E-09 | 0.12 |
| Hip circumference | 25673412 | 0.21 | 0.043 | 4.91 | 8.89E-07 | 0.13 |
| Obesity class 1 | 23563607 | 0.23 | 0.047 | 4.82 | 1.44E-06 | 0.22 |
| Waist-to-hip ratio | 25673412 | 0.19 | 0.042 | 4.49 | 7.10E-06 | 0.11 |
| Obesity class 2 | 23563607 | 0.26 | 0.059 | 4.33 | 1.46E-05 | 0.19 |
| Obesity class 3 | 23563607 | 0.31 | 0.075 | 4.09 | 4.34E-05 | 0.13 |
| Body mass index | 20935630 | 0.17 | 0.045 | 3.90 | 9.63E-05 | 0.19 |
| Overweight | 23563607 | 0.18 | 0.048 | 3.72 | 0.0002 | 0.11 |
| Extreme bmi | 23563607 | 0.20 | 0.074 | 2.64 | 0.008 | 0.69 |
| Height_2010 | 20881960 | 0.07 | 0.041 | 1.66 | 0.10 | 0.29 |
| Extreme height | 23563607 | 0.09 | 0.059 | 1.59 | 0.11 | 1.27 |
| Extreme waist-to-hip ratio | 23563607 | 0.06 | 0.095 | 0.66 | 0.51 | 0.36 |
| **Depression-related phenotypes** |  |  |  |  |  |  |
| Depressive symptoms | 27089181 | 0.52 | 0.060 | 8.66 | 4.54E-18 | 0.05 |
| Self-reported depression | UK<br>biobank | 0.46 | 0.072 | 6.43 | 1.27E-10 | 0.02 |
| Major depressive disorder | 22472876 | 0.48 | 0.089 | 5.37 | 7.67E-08 | 0.17 |
| **Arthritis-related phenotypes** |  |  |  |  |  |  |
| Self-reported osteoarthritis | UK<br>biobank | 0.63 | 0.066 | 9.62 | 6.82E-22 | 0.02 |
| Osteoarthritis of the knee (using ICD10 diagnostic codes) | UK<br>biobank | 0.49 | 0.085 | 5.75 | 8.67E-09 | 0.01 |
| Osteoarthritis of the hip (using ICD10 diagnostic codes) | UK<br>biobank | 0.17 | 0.095 | 1.77 | 0.08 | 0.01 |
| Self-reported arthritis (not otherwise specified) | UK<br>biobank | 0.58 | 0.316 | 1.84 | 0.07 | 0.00 |
| Rheumatoid arthritis | 24390342 | 0.11 | 0.057 | 1.96 | 0.0497 | 0.16 |

Nominally significant genetic correlations **in bold**
\*All GWAS results in participants of European ancestry; SNPs from the MHC (chr6 26M~34M) region was removed for all traits
<sup>a</sup>genetic correlations with chronic back pain (results from the discovery stage meta-analysis)

Bjornsdottir G, Benonisdottir S, Sveinbjornsson G, et al. Sequence variant at 8q24.21 associates with sciatica caused by lumbar disc herniation. Nat Commun 2017;8:14265. doi: 10.1038/ncomms14265

**Supplemental Figure S1.**
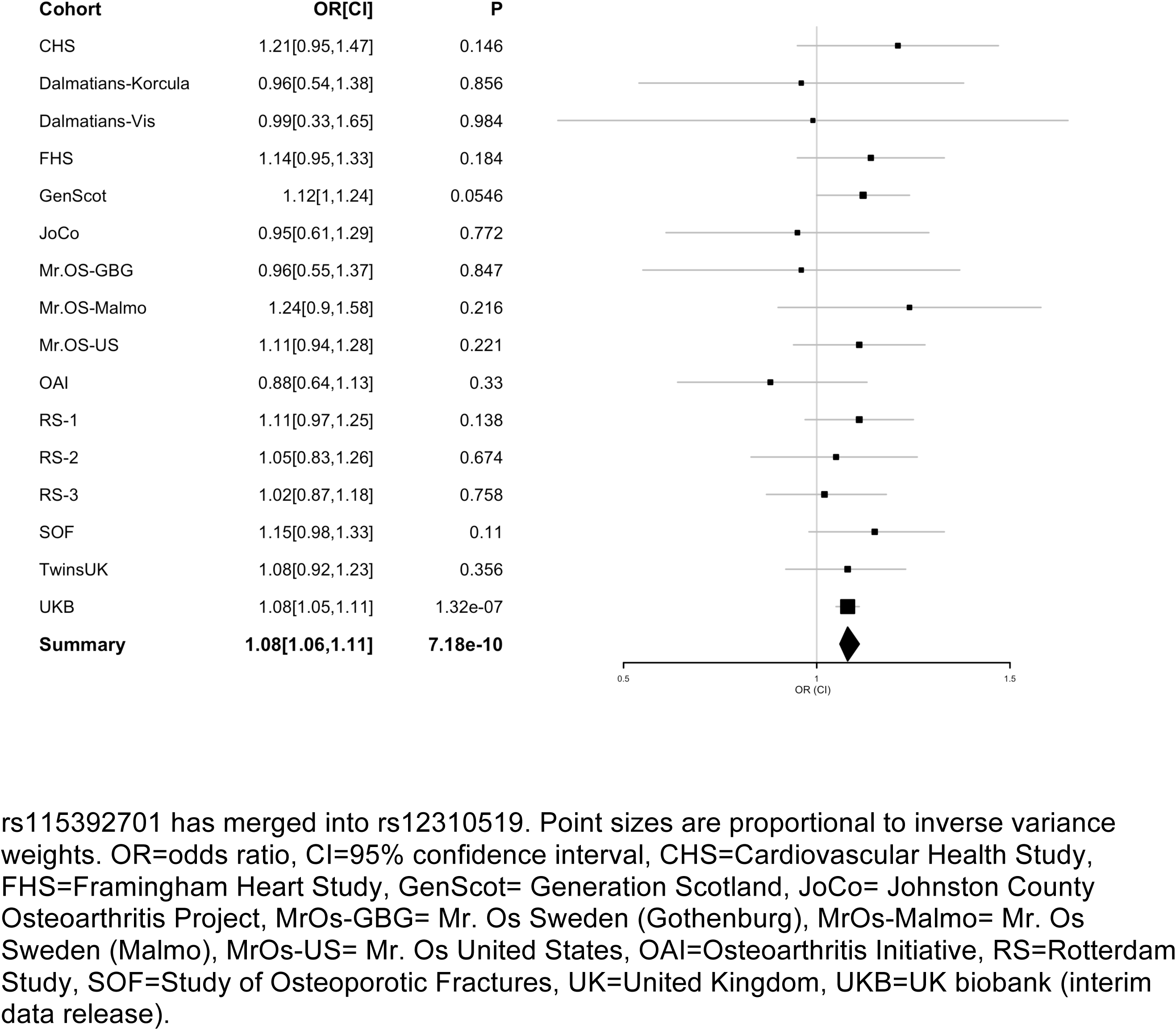
Forest plot of rs12310519 (*SOX5*, chr12) association with chronic back pain in the meta-analysis (discovery)

**Supplemental Figure S2.**
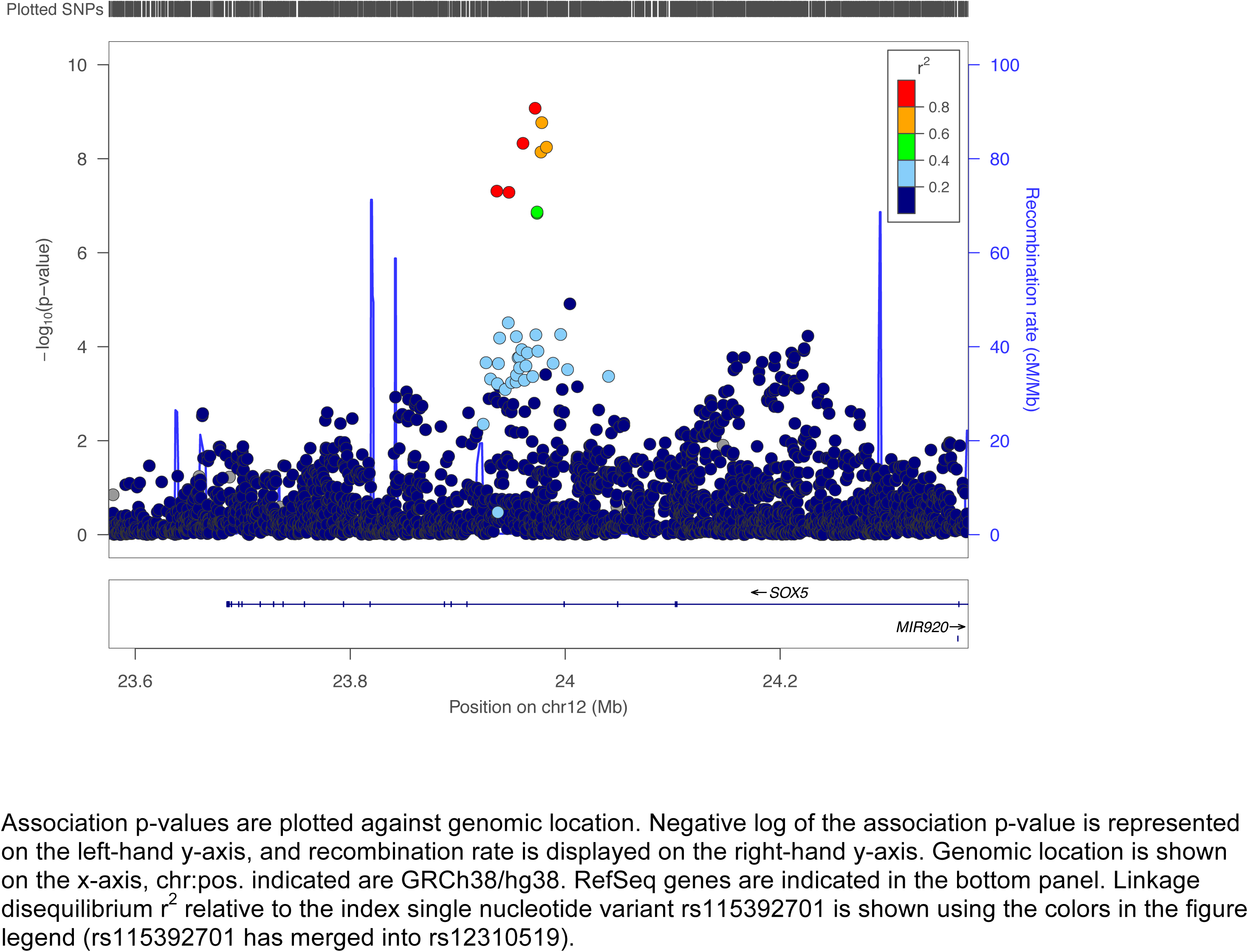
Regional association plot of *SOX5* locus

**Supplemental Figure S3.**
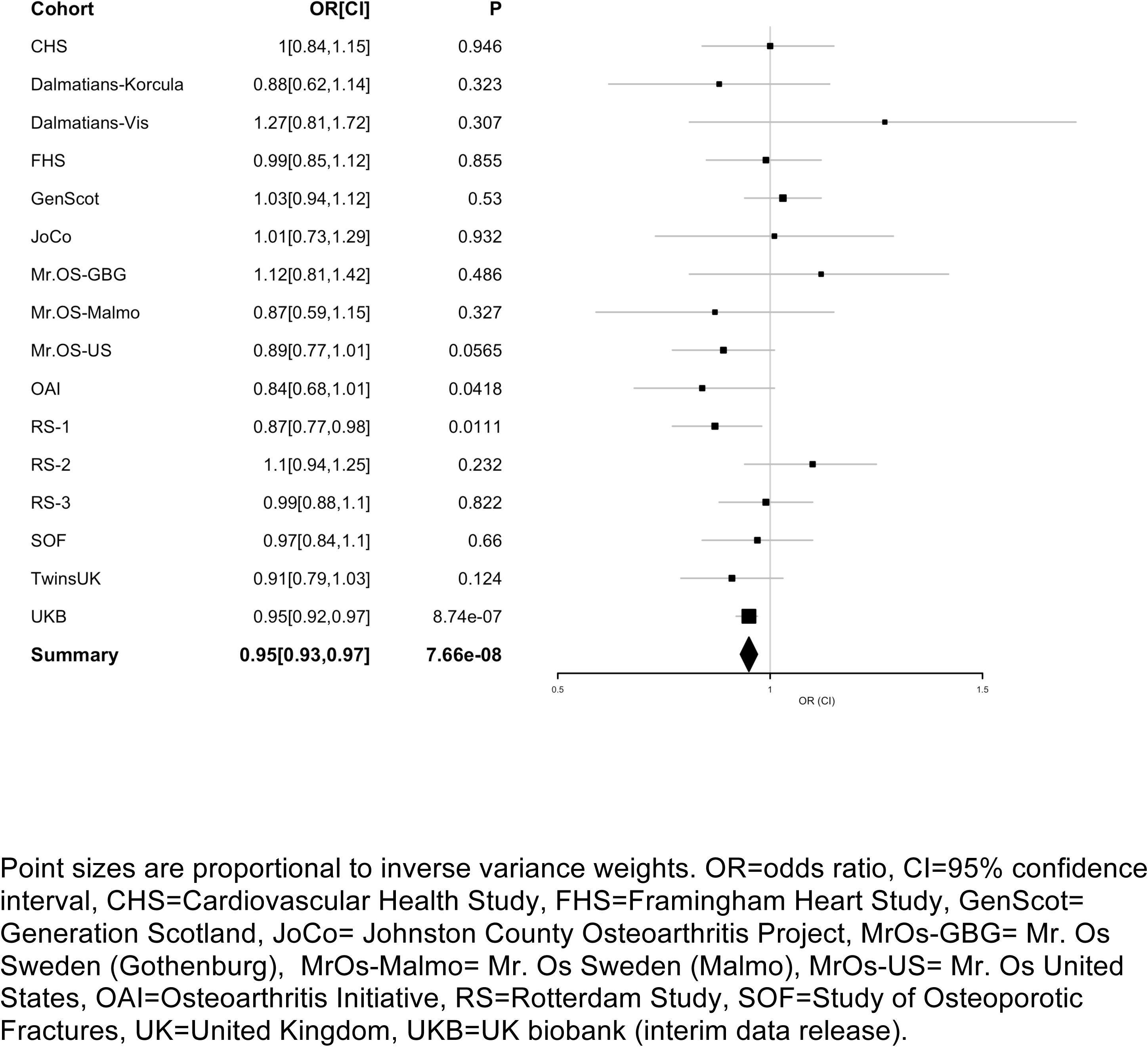
Forest plot of rs1453867 (*DIS3L2,* chr2) association with chronic back pain in the meta-analysis (discovery)

**Supplemental Figure S4.**
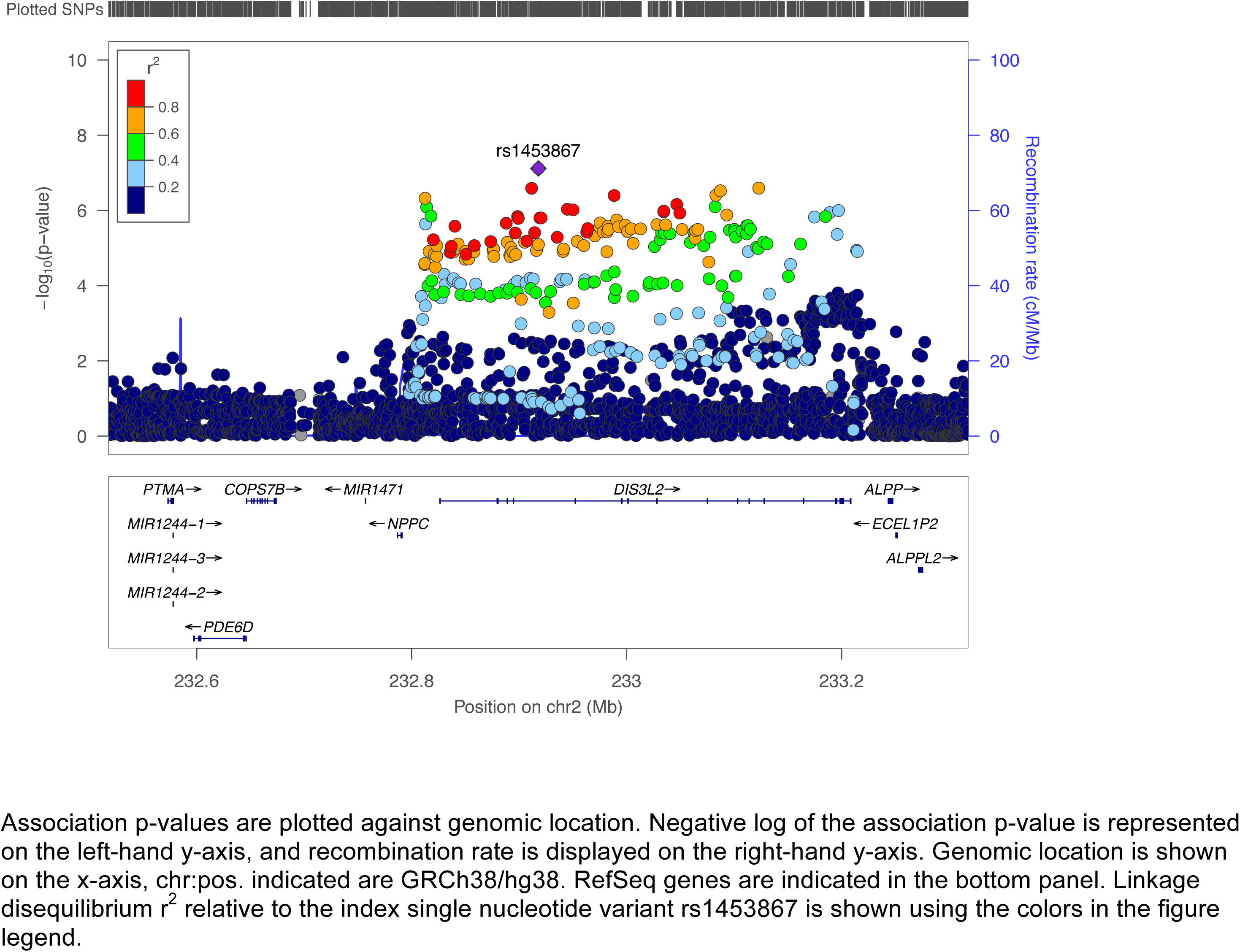
Regional association plot of *DIS3L2* locus

**Supplemental Figure S5.**
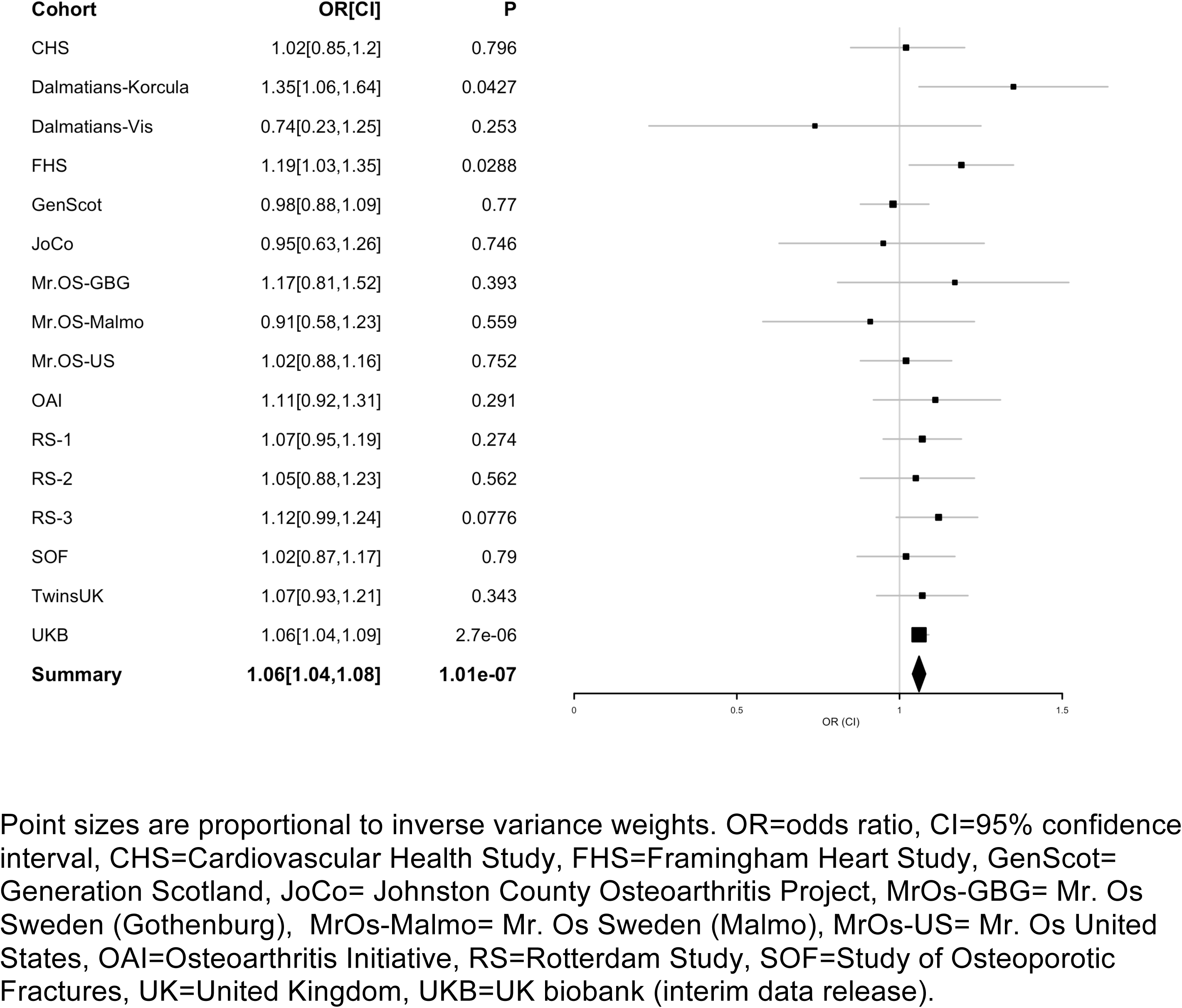
Forest plot of rs7833174 (*CCDC26/GSDMC,* chr8) association with chronic back pain in the meta-analysis (discovery)

**Supplemental Figure S6.**
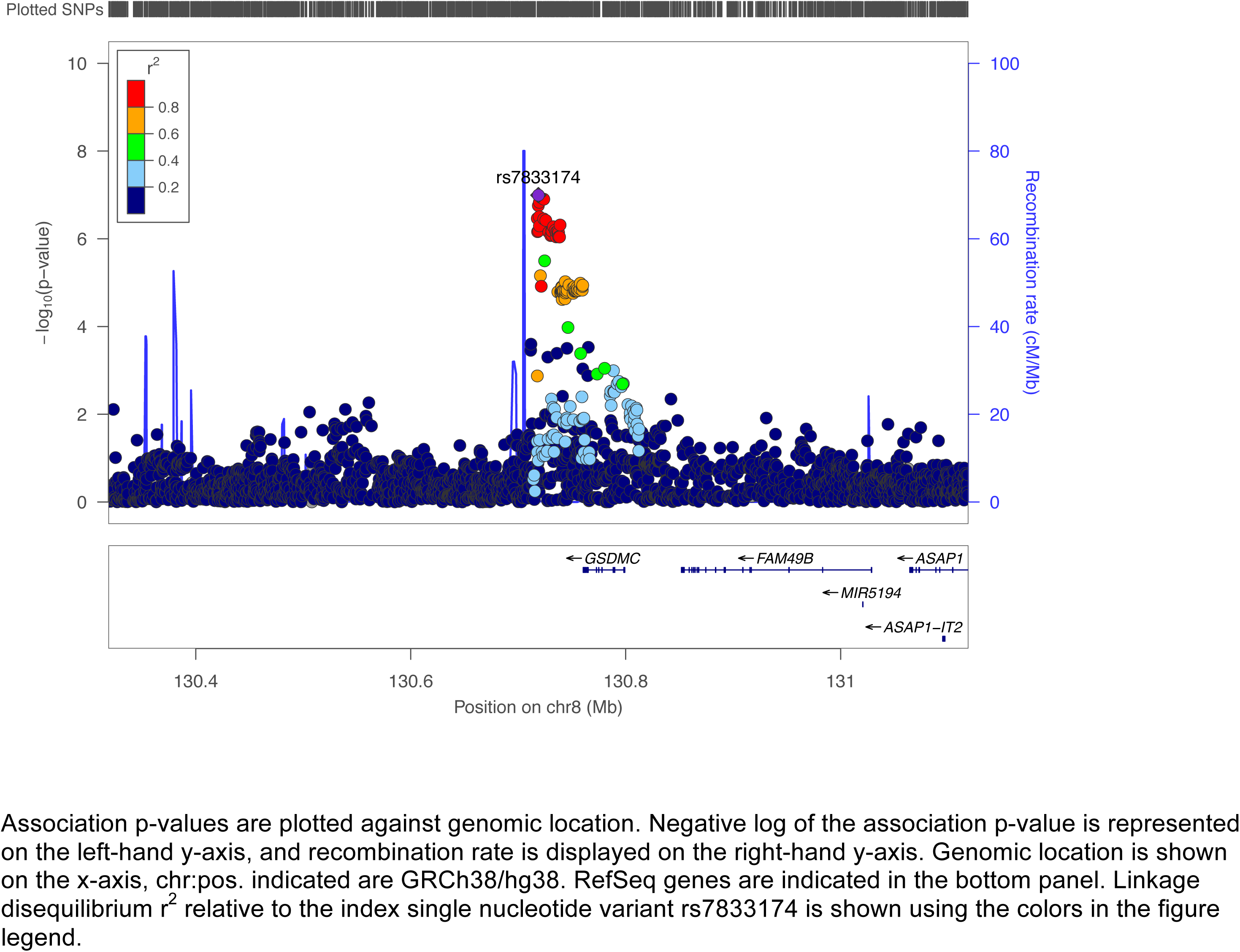
Regional association plot of *CCDC26/GSDMC* locus

**Supplemental Figure S7.**
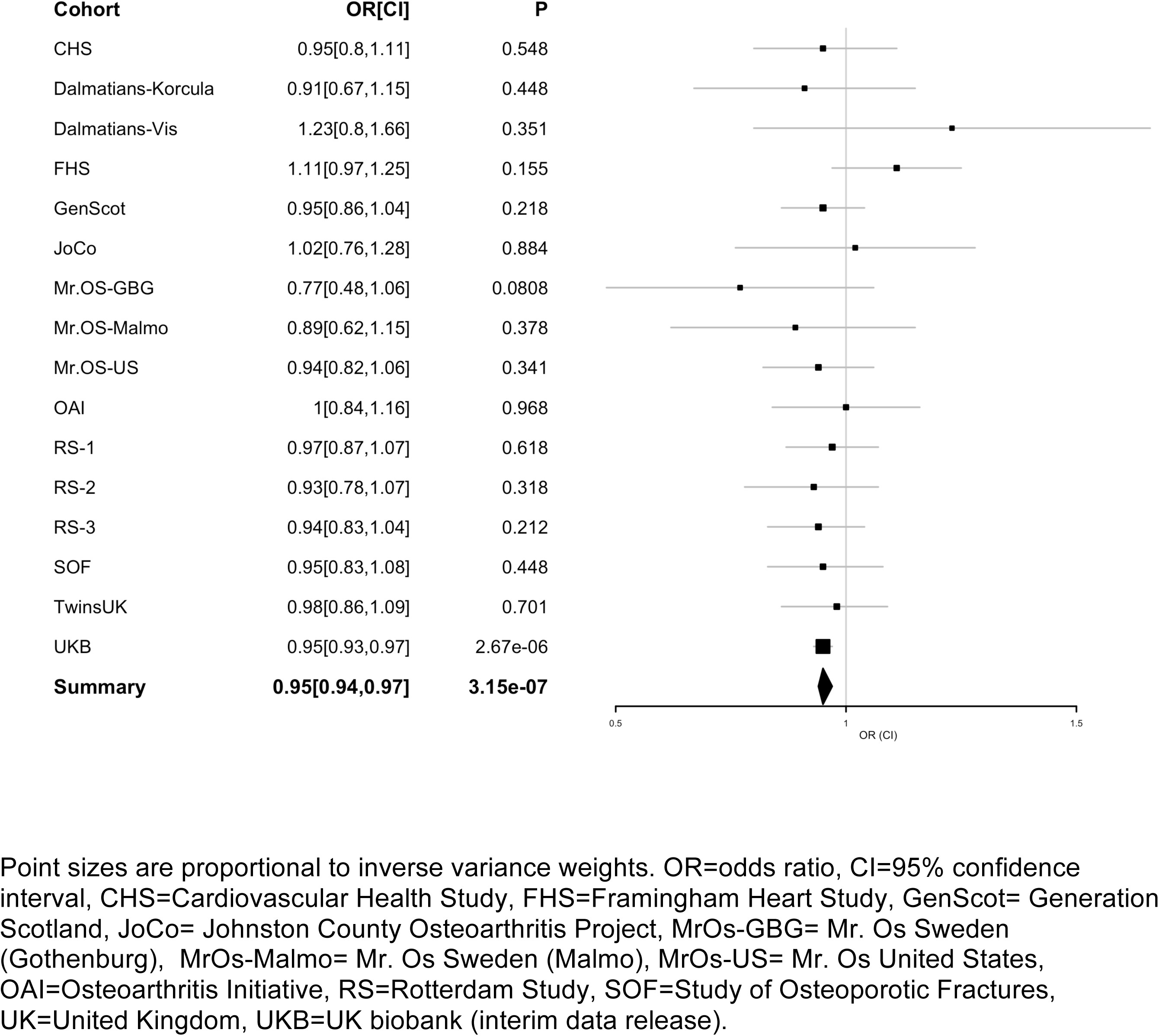
Forest plot of rs4384683 (*DCC,* chr18) association with chronic back pain in the meta-analysis (discovery)

**Supplemental Figure S8.**
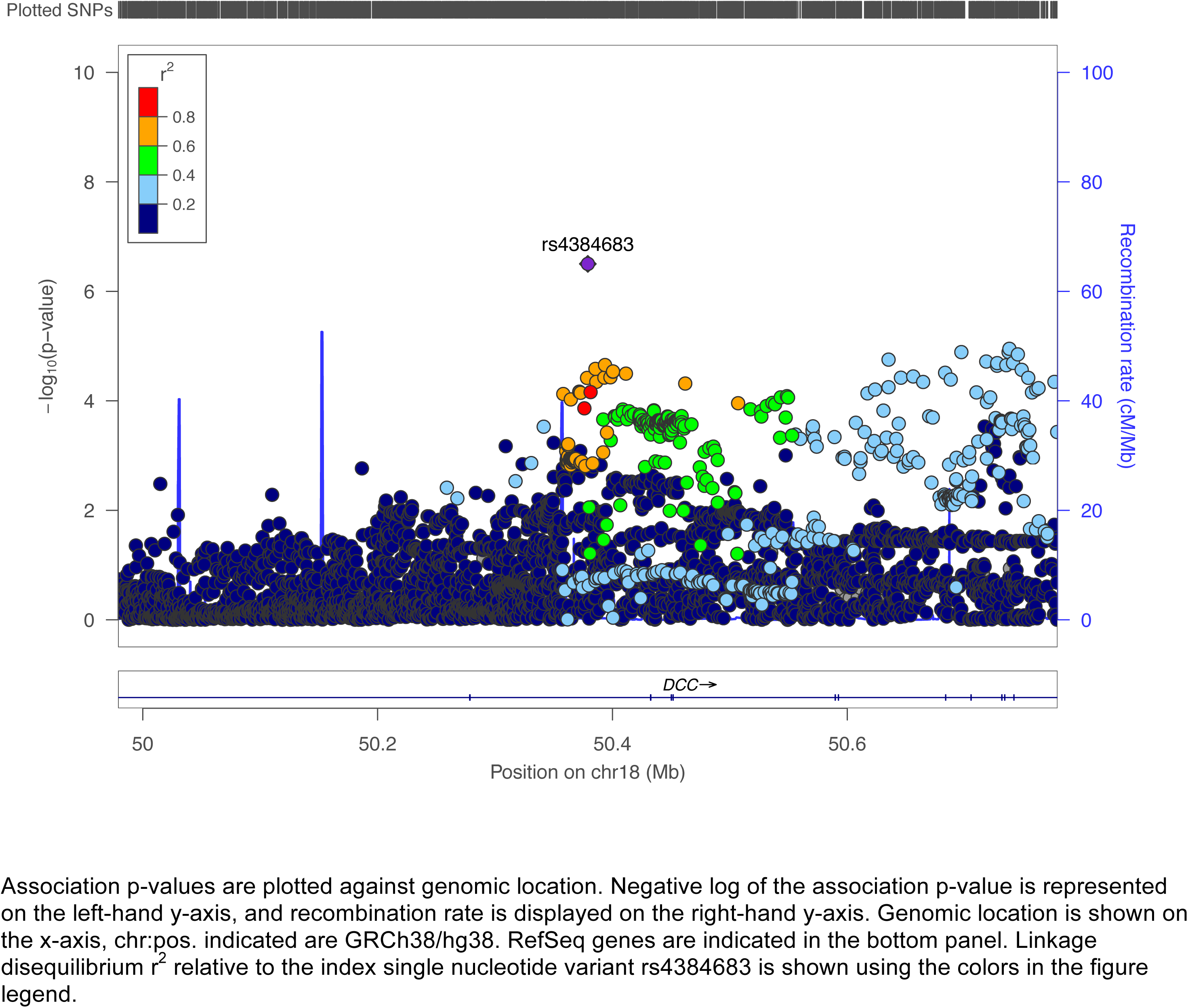
Regional association plot of *DCC* locus

## Supplemental Methods

### Cohort Descriptions and Cohort-Specific Phenotype Descriptions

See Supplemental Table S2 for further details of chronic back pain phenotype definitions for each cohort.

**Cardiovascular Health Study (CHS)**- CHS is a community-based observational study of risk factors for cardiovascular disease in ambulatory adults 65 years or older.^1^ Participants were recruited from Medicare enrollment lists in four U.S. communities: Forsyth County, North Carolina; Sacramento County, California; Washington County, Maryland; and Pittsburgh, Pennsylvania. Starting in 1989, 5201 participants underwent extensive annual clinical examinations. An additional predominantly African-American cohort of 687 individuals was enrolled in 1992-1993. Study measures included traditional cardiovascular risk factors such as blood pressure and lipids as well as objective measures of subclinical disease, including echocardiography of the heart, carotid ultrasound, and cranial magnetic-resonance imaging. African-Americans were not included in the current study due to the meta-analysis’ focus on individuals of European ancestry only. All participants gave informed consent to use genetic data for analyses. A variety of self-report measures were also collected. Back pain questions began during the 2^nd^ examination, and included the question “In the past year, have you had pain in your back for more than half the days of any month?”.

**Framingham Heart Study (FHS)-** FHS began in 1948 as a longitudinal population-based cohort study of the causes of heart disease.^2^ The Original cohort was comprised of 5,209 men and women from the town of Framingham, Massachusetts. In 1971, 5,124 offspring of the original Framingham cohort and their spouses were entered in the Offspring cohort.^3^ Examination 8 of the Framingham Offspring cohort study began in 2005 and concluded in 2008. Participants were asked “Have you had back pain in the past 12 months?”, with response options including “no”, “a few days”, “some days”, “most of the days”, and “all days”.

**Generation Scotland (GS)-** GS is a family-based genetic epidemiology study with genetic, sociodemographic, and clinical data from approximately 24,000 volunteers across Scotland aged 18–98 years.^4^ Participants were recruited through a variety of methods from February 2006 to March 2011. Biological samples and anonymized data form a resource for research on the genetics of health, disease and quantitative traits of current and projected public health importance. A ‘broad’ form of consent was acquired from participants to use their data and samples for a wide range of medical research, including re-contact for the potential collection of other data or samples, or for participation in related studies.^4^ Participants reporting any musculoskeletal pain were asked whether specific areas of pain or discomfort had been going on for more than 3 months. Participants specified ‘yes’ or ‘no’ for back pain of duration > 3 months.

**Johnston County Osteoarthritis Project (JoCo)-** JoCo is an ongoing, population-based prospective cohort begun in 1990 to fill knowledge gaps about prevalence, incidence, and progression of OA, and its risk factors, in Caucasian and African-American men and women in North Carolina.^5^ Participants were recruited from six townships and surrounding rural areas, and eligible individuals completed two home interviews and clinic examinations. The sample has been followed approximately every 5 years since inception with repeated surveys, radiographic, and clinical assessments, and other diagnostic testing. Beginning in 2012, participants were asked a series of questions about areas of pain experienced in the last 12 months, including middle back (thoracic) and/or lower back (L1 through S1) pain that they had experienced on most days of any one month in the last year. Those who reported pain in the middle back or lower back were then asked the number of months in the past year during which they had had pain in this location on most days of the month.

**MrOs Sweden-** MrOs Sweden is a population-based cohort study of community-living Swedish men aged 69–81, part of a multi-national collaboration including the MrOs US study and MrOs Hong Kong.^6^ MrOs Sweden includes 3,014 men recruited from 3 study centers, including Gothenburg, Malmo, and Uppsala. Participants were randomly selected from the Swedish national population register. At the study baseline, participants answered a questionnaire including fracture history, lifestyle, and clinical symptoms.^6^ Participants were asked “Have you had back pain in the past 12 months?”. Response options included “no”, “never”, “rarely”, “some of the time”, “most of the time”, and “all of the time”. Only participants from the Gothenburg and Malmo sites who had available genetic and back pain data were included in the current study.

**MrOs US (The Osteoporotic Fractures in Men)** - MrOs US is a prospective, population-based cohort study of older men recruited from the geographic regions surrounding Birmingham, Minneapolis, Palo Alto, Pittsburgh, Portland, and San Diego. The study was designed to determine the extent to which fracture risk is related to bone mass and geometry, lifestyle, anthropometric, and neuromuscular measures, as well as to determine how fractures affect quality of life in older men.^7^ MrOs US is part of a multi-national collaboration including the MrOs Sweden study and MrOs Hong Kong. At baseline, MrOs participants completed questionnaires regarding medical history, medications, physical activity, diet, alcohol intake, and cigarette smoking. Objective measures of anthropometric, neuromuscular, vision, strength, and cognitive variables were also obtained. MrOs participants were asked the question “During the past 12 months have you experienced any back pain?”, with response options including “no”, “never”, “rarely”, “some of the time”, “most of the time”, and “all of the time”.

**Osteoarthritis Initiative (OAI)**- The OAI is a multicenter, longitudinal, prospective observational study of knee osteoarthritis. It is a public-private partnership between the NIH and private industry that seeks to develop a public domain research resource to facilitate the scientific evaluation of biomarkers for osteoarthritis as potential surrogate endpoints for disease onset and progression.^8^ OAI includes 4,796 participants age 45-79 recruited from clinical centers in Maryland, Ohio, Pennsylvania, and Rhode Island. Participants completed serial assessments and examinations annually since study inception, with a smaller subset seen more frequently for analysis of change over shorter intervals. At each annual assessment, participants were asked the question ““How often were you bothered by back pain in the past 30 days?”, with response options including “no”, “rarely”, “some of the time”, “most of the time”, and “all of the time”.

**Rotterdam Study (RS)-** Rotterdam Study is a prospective population-based cohort study on the determinants of chronic disabling diseases, ongoing since 1990.^9^ The first Rotterdam cohort (RS-1) included 7,983 persons ≥55 years of age living in the Ommoord district in the city of Rotterdam, in the Netherlands. In 2000, 3011 additional participants ≥55 years of age were added to the cohort (RS-2). In 2006, a further extension of the cohort was initiated in which 3,932 subjects living in the Ommoord district were included, aged 45–54 years (RS-3)^9^. Back pain was assessed at several of the RS examinations. In RS-1, back pain was assessed with the question, “Have you had pain in the low back in the last month?”, with further questions as to the duration of pain in the low back. In RS-2 and RS-3, back pain was assessed with the question, “Do you have pain or stiffness in your back?”, with further questions as to the duration of pain in the back.

**Study of Osteoporotic Fractures (SOF)**- SOF is a community-based multi-center prospective observational study of a cohort of 10,366 women ≥55 years of age who were recruited from four metropolitan areas in the US: Minneapolis, Pittsburgh, Portland, and Baltimore. The primary purpose of SOF was to describe the risk factors for osteoporotic fractures in women.^10^ Participants were initially recruited from mailings to age-eligible women identified from community-based listings, such as memberships of large health maintenance organizations. SOF is funded by NIA. SOF participants were asked the question “During the past 12 months have you experienced any back pain?”, with response options including “no”, “never”, “rarely”, “some of the time”, “most of the time”, and “all of the time, constantly”.

**10,001 Dalmatians-** This study aims to identify genetic and environmental determinants of health and disease in genetic-isolate island populations from Dalmatia, Croatia. The Vis and Korcula cohorts included unselected Croatians who were recruited from villages on the Dalmatian islands of Vis and Korcula. Participants completed questionnaires regarding lifestyle and environmental exposures, biochemical and physiological measurements, and genotyping. Participants were asked the questions “Have you ever had an episode of chronic LBP that lasted for over 3 months?” and “Do you have low back pain at the moment?”, and participants who responded yes to both questions were considered to have chronic back pain of duration > 3 months.

**TwinsUK**- TwinsUK is a British adult twin registry originally assembled to study the heritability and genetics of age-related diseases^11^. The registry was started in 1992 and consists of approximately 13,000 monozygotic and dizygotic adult Northern European twins aged 16 to 85 years from all over the UK, as well as some parents and siblings. Individuals were recruited from the general population through national media campaigns throughout the UK. TwinsUK is generally representative of singletons and the United Kingdom population. TwinsUK participants were asked the question “In the past 3 months, have you had pain in your back on most days?”. Further questions inquired about back pain in specific locations while referencing a mannekin showing locations of the pain in the thoracic and lumbar regions, and whether any such pain had persisted for at least the past 3 months.

**UK Biobank (UKB)-** The UK (United Kingdom) Biobank is a population-based prospective study involving more than 500,000 participants, established to allow detailed studies of the genetic and nongenetic determinants of the diseases of middle age and old age.^12^ UKB aims to combine extensive assessment of exposures with comprehensive follow-up and characterization of many different diseases and health-related outcomes, as well as to promote innovative science by maximizing access to the resource. The study thus far has collected and continues to collect longitudinal data and extensive phenotypic and genotypic detail about its participants, including data from questionnaires, physical measures, diagnostic imaging, and genotyping. Participants reported locations (including back pain) where they had experienced pain in the last month that interfered with usual activities. Participants reporting back pain specified ‘yes’ or ‘no’ for back pain of duration > 3 months.

### Statistical Analysis (Detailed Description)

Discovery meta-analysis included adults of European ancestry from these 16 population- and community-based cohorts listed above. Replication was conducted in an independent sample of UKB European ancestry participants not included in the UKB interim data release that was used for discovery, and a joint (discovery-replication) meta-analysis was performed

For the discovery stage, we conducted genome-wide association analyses in each of the 16 cohorts, and subsequent meta-analysis of autosomal SNPs to combine results from all cohorts. Each site conducted GWAS using logistic regression models with additive genetic effects to test for associations between each variant and chronic back pain as a binary trait. These models adjusted for age, sex, and study-specific covariates such as study site, clinic site, or array. (Supplemental Table S3) Each cohort evaluated population substructure and adjusted for principle components where indicated (Supplemental Table S3). Height and body mass index (BMI), calculated as weight in kilograms divided by height in meters squared, were not included as covariates in site-specific GWAS, since these traits might lie along the causal pathway or (in the case of BMI) reflect a consequence of CBP. Harmonization and quality control for GWAS results from each cohort were conducted using the EasyQC software package in the R statistical environment (v3.2.2), using methods described previously.^13^ After removal of SNPs with low minor allele frequencies (<0.005 for UKB, <0.03 for Vis, <0.01 for other cohorts) or imputation quality (<0.7 for UKB, <0.6 for other cohorts), deviation from Hardy-Weinberg equilibrium (p < 1 × 10-6), low number of cases (<15) or controls (<15), large absolute values of beta coefficients (≥10), and low minor allele count (≤10), call rates <0.95, the range of SNPs included in the meta-analysis was between 6,205,227 (Croatia-Vis) and 9,775,703 (MrOs-Gothenburg) (Supplemental Table S3). Fixed-effect inverse-variance weighted meta-analysis was performed using METAL (http://csg.sph.umich.edu/abecasis/metal/). Quality control and meta-analysis were conducted twice, independently of each other, by researchers at the University of Washington (MP and PS) and PainOmics (YA, LCK, and YT). The results from the two centers were compared to ensure accuracy. We used linkage disequilibrium (LD) score regression (LDsr) to examine potential confounding.^14^ The genome-wide significance level was defined as p<5×10^−8^, and suggestive significance level was defined as p<5×10^−7^, after using the LDsr intercept as a correction factor. Q-Q and Manhattan plots were generated in R. Effect heterogenity across studies was quantified using the I^2^ metric (range 0-100%).^15^ Between-study heterogeneity was tested using the Cochran Q statistic which was considered significant at p<0.1. We used LocusZoom (http://csg.sph.umich.edu/locuszoom/) to evaluate regional association plots at regions of interest, and conducted Genome-Wide Complex Trait Analysis conditional and joint analyses (http://cnsgenomics.com/software/gcta/#About) for variants in each locus exceeding the suggestive significance level, conditional on the most significant variant at each locus.

The most highly associated at genome-wide significant loci were subjected to replication in the independent sample of UKB participants. Analysis in the independent sample used logistic regression with additive genetic effects, adjusting for age, sex, array, and 10 principal components (Supplemental Table S3); significant replication was defined using a Bonferroni-corrected threshold of p<0.05 divided by the number of genome-wide significant loci (0.05/1). The most highly-associated variants at loci with suggestive significance were selected for a joint (discovery-replication) meta-analysis using p<5×10–8 to define genome-wide significance.

For genome-wide significant variants and those in LD (r^2^ >0.6) we examined previously reported GWAS associations with other phenotypes using PhenoScanner v1.1 (http://www.phenoscanner.medschl.cam.ac.uk/phenoscanner), and conducted functional annotation using FUMA (http://fuma.ctglab.nl). FUMA draws upon multiple publicly available databases, annotating variants for functional consequences on gene functions using the combined annotation dependent depletion (CADD) score,^16^ potential regulatory functions (RegulomeDB score),^17^ and effects on gene expression using expression quantitative trait loci (eQTLs) of different tissue types (GTExv6 and other databases for eQTLs)^18,19^ (Supplemental File). The CADD score is computed based on 63 annotations; the higher the score, the more potentially deleterious the variant is. A CADD score of ≥10 indicates a variant predicted to be among the 10% most deleterious substitutions involving the human genome, a score of ≥20 indicates the 1% most deleterious, and soforth. We used data from the Roadmap Epigenomics Project to evaluate whether the lead variants at each locus reside in enhancer regions for tissue types of interest by comparing CHIP-seq signals for chromatin state markers in selected tissues with possible conceptual connections to back pain via roles involving chondrogenesis, vertebral development, muscle, and the central nervous system.^20,21^

Because genome-wide significant variants at *SOX5* and *CCDC26/GSDMC* were found to be associated with height in prior published GWAS, we conducted *post hoc* region-specific GWAS secondary analyses accounting for possible interrelationships with height. Model 1 (CBP ~ SNP + age + sex + array + PC1 + … + PC10) examined associations between SNPs and CBP using logistic regression, among UKB participants from the interim data release. Model 2 (CBP ~ SNP + age + sex + + height+ array + PC1 + … + PC10) used similar methods, but also adjusted for height. Models 1 and 2 were compared descriptively with respect to the lead SNPs in *SOX5* and *CCDC26/GSDMC* that were identified in the discovery stage meta-analysis, to assess whether these SNP-CBP associations differed when including height as a covariate. Model 3 (CBP ~ SNP + age + sex + array + PC1 + … + PC10) was the same as Model 1, but analysis was conditional on the top height-associated variant in the region. Models 1 and 3 were compared descriptively with respect to the lead SNPs from region-specific analyses among participants from the UKB interim data release, to assess whether results for the lead SNPs differed when conditional on the top height-associated variant in the region. Model 4 was a linear regression model with additive genetic effects to examine for associations between each variant and height (height ~ SNP + age + sex + array + PC1 + … + PC10). Model 5 was the same as Model 4 (height ~ SNP + age + sex + array + PC1 + … + PC10), but analysis was conditional on the top CBP-associated variant in the region. Models 4 and 5 were compared descriptively with respect to the lead SNPs from region-specific analyses among participants from the UKB interim data release, to assess whether results for the lead SNPs differed when conditional on the top CBP-associated variant in the region (Supplemental Table S7).

Last, we used LDsr of summary-level GWAS results from the discovery stage to estimate heritability due to common autosomal SNPs and genetic correlations.^14^ We transformed the observed SNP heritability to the liability scale, in order to make heritability estimates for CBP comparable with traditional heritability estimates from twin studies.^22^ We used cross-trait LDsr and publicly available meta-GWAS results from LDhub to examine genetic correlations with a limited list of traits with strong conceptual links to CBP, including anthropometrics (adult height, infant/childhood height, waist/hip circumference, BMI, and overweight/obesity), depression and depressive symptoms, osteoarthritis and rheumatoid arthritis.^23,24^

---

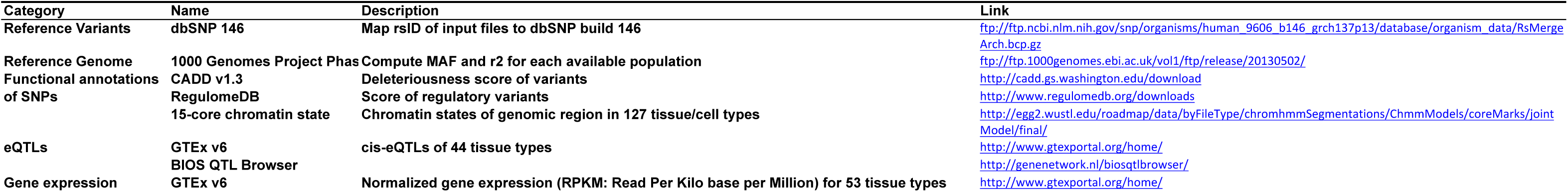
List of data repositories and tools used in FUMA in the current study.

**Table.1.**
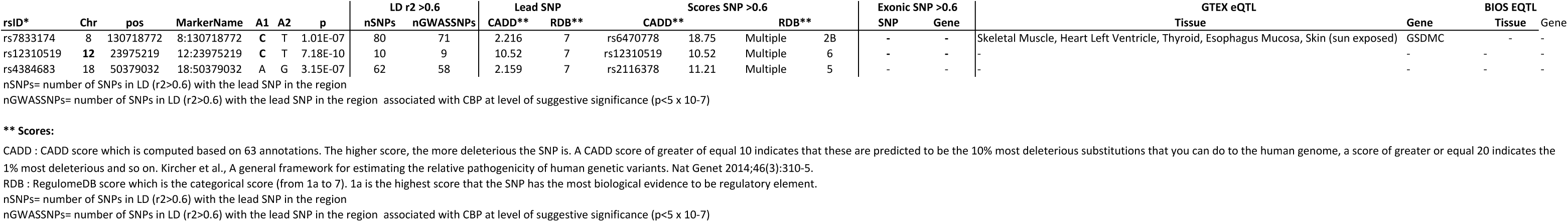
Summary of FUMA results.

**Table2.** Lead SNPs.

| No | GenomicLocus | uniglID | rsID | chr | pos | p | nIndSigSNPs | IndSigSNPs |
| --- | --- | --- | --- | --- | --- | --- | --- | --- |
| 1 | 1 | 8:130718772:C:T | rs7833174 | 8 | ##### | 1.01E-07 | 1 | rs7833174 |
| 2 | 2 | 12:23975219:C:T | rs12310519 | 12 | 23975219 | 7.18E-10 | 2 | rs12310519;rs12308843 |
| 3 | 3 | 18:50379032:A:G | rs4384683 | 18 | 50379032 | 3.15E-07 | 1 | rs4384683 |

nIndSignSNPs= number of 'independent' significant SNPs (independent SNPs refers to those not in LD with the lead SNP; it does not refer to independence conditional on the lead SNV using GCTA COJO analyses, as described in the manuscript).

**Table3.**
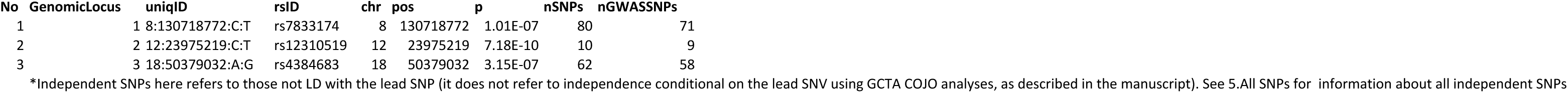
Independent SNPs (r2 >0.6)*

**Table4.**
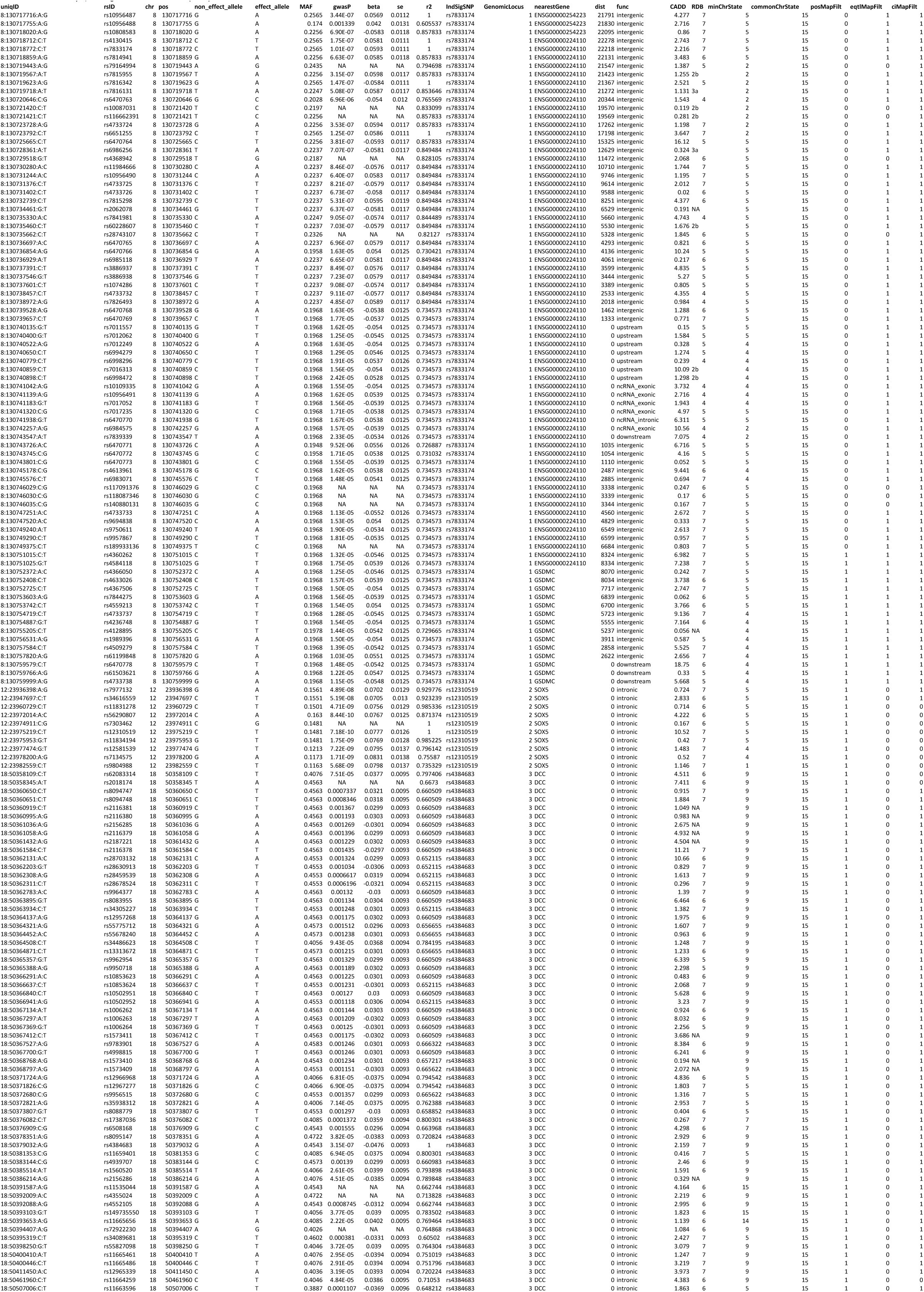
SNPs in FUMA Analysis, including Lead SNPs and those in LD (r2>0.6)

**Table5.**
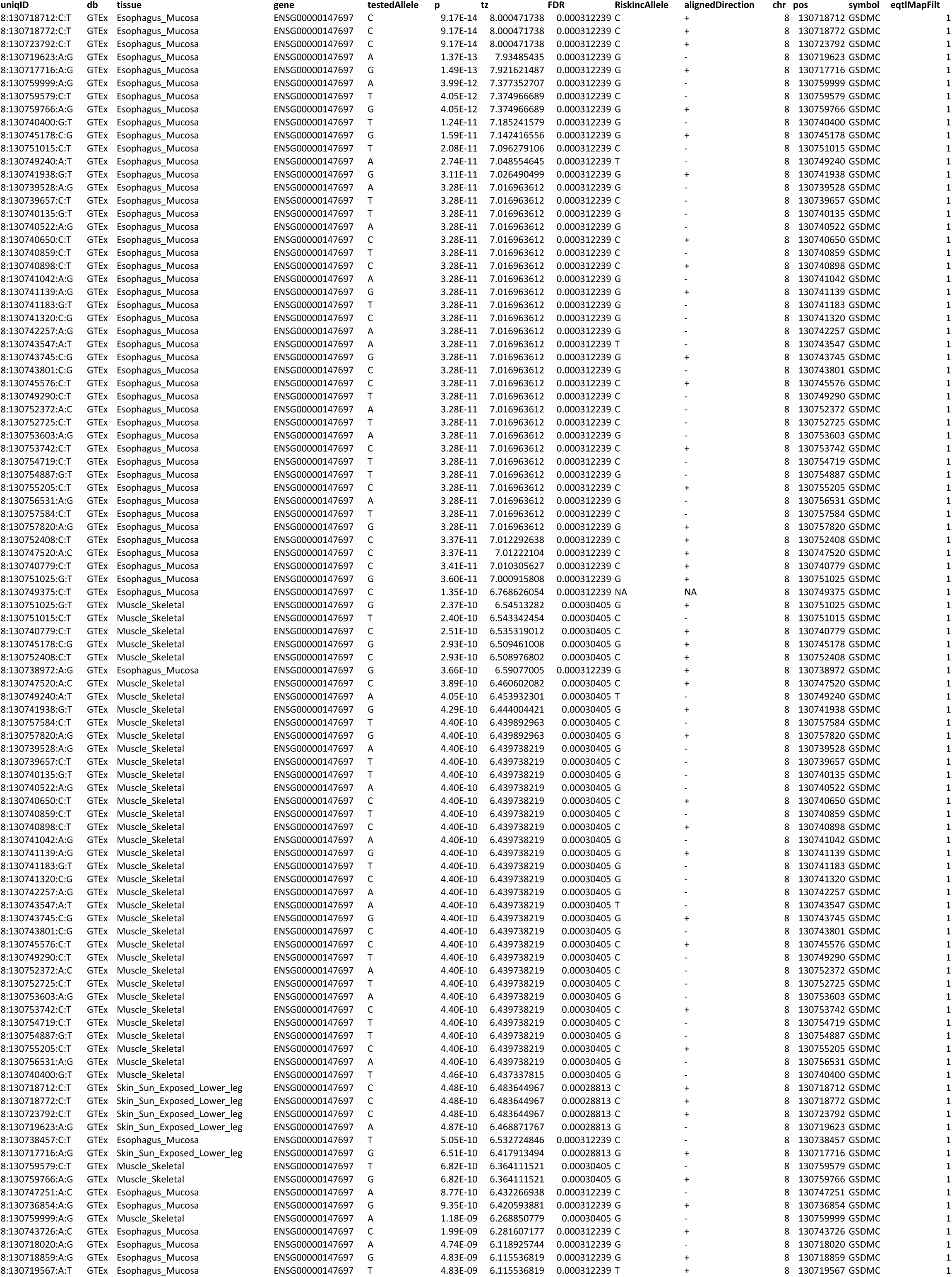
eQTL.results (SNPs, LD r2>0.6)

**Table5.**
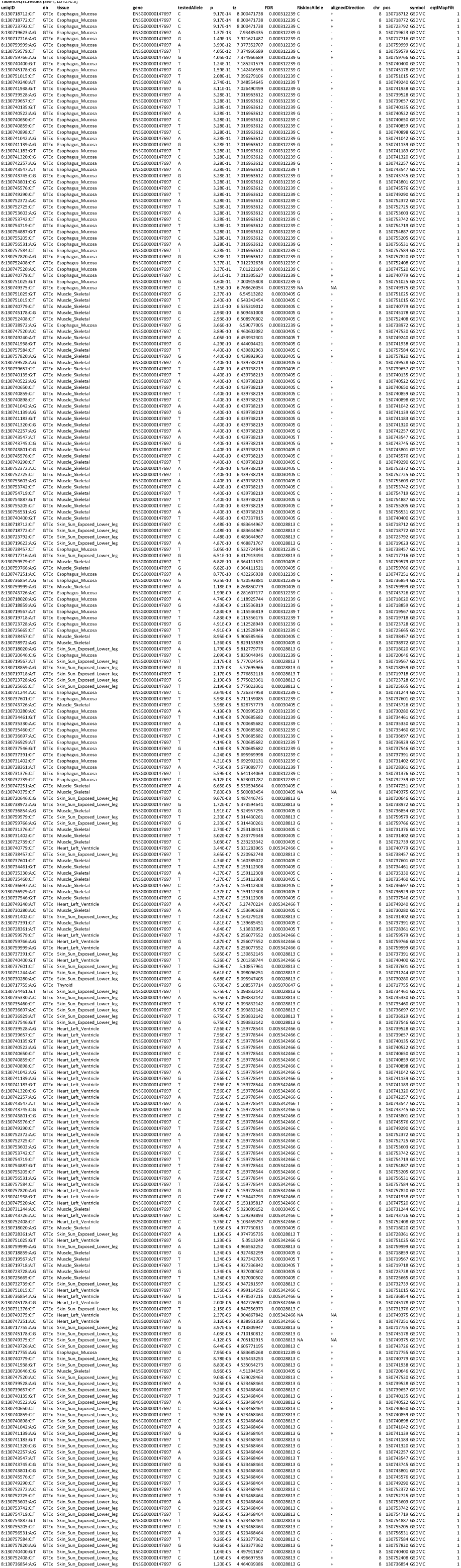
eQTL.results (SNPs, LD r2>0.6)

**Table6.**
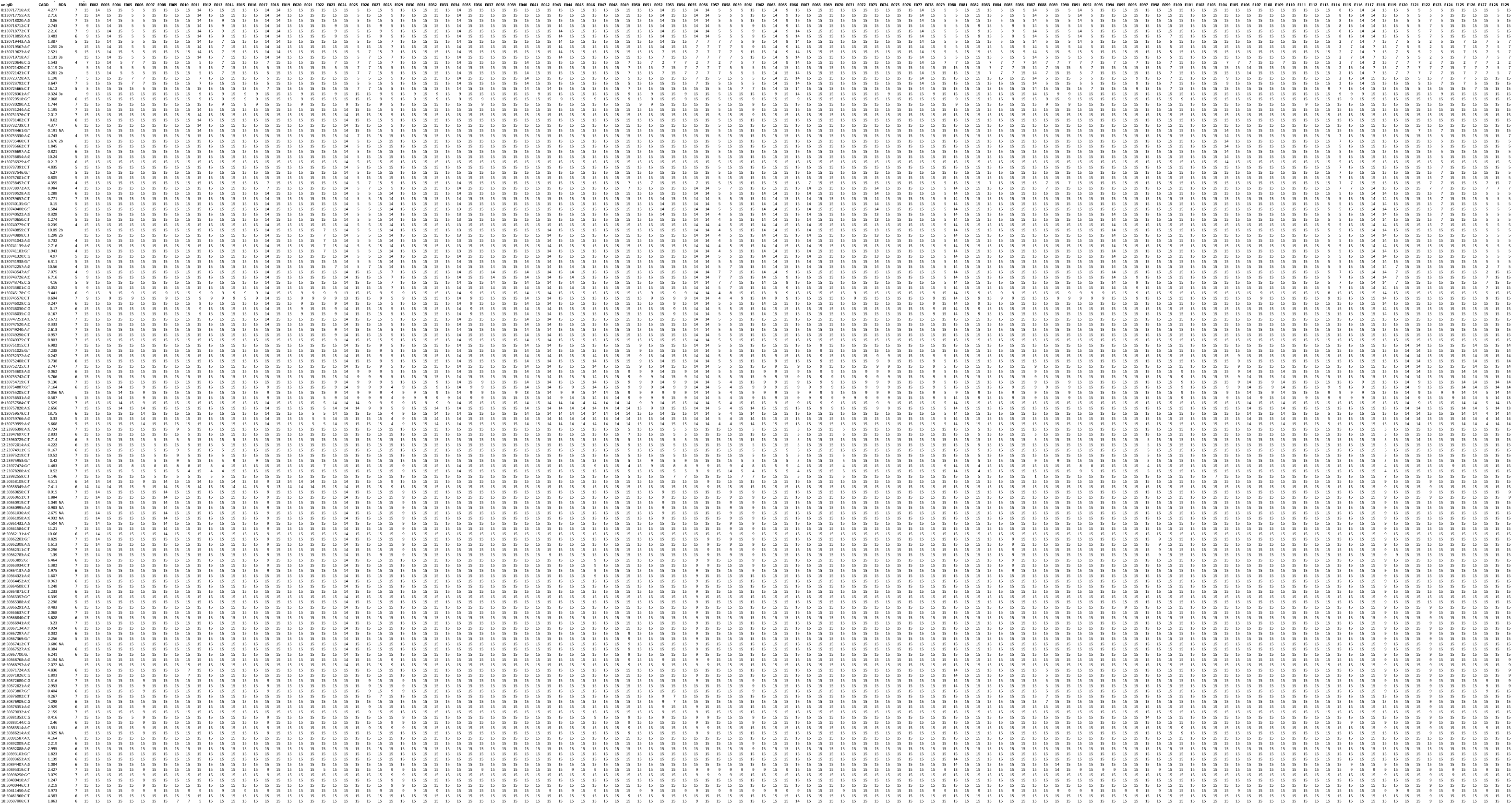
Functional annotation (SNPs LD r2>0.06), E nr = ROADMAP identifier.

